# The association of DNA methylation with body mass index: distinguishing between predictors and biomarkers

**DOI:** 10.1101/2019.12.16.877464

**Authors:** Zoe E. Reed, Matthew J. Suderman, Caroline L. Relton, Oliver S.P. Davis, Gibran Hemani

**Affiliations:** Medical Research Council Integrative Epidemiology Unit (MRC IEU), Department of Population Health Sciences, Bristol Medical School, University of Bristol, UK; The Alan Turing Institute, British Library, 96 Euston Road, London, UK

**Keywords:** BMI, DNA methylation, ALSPAC, ARIES, Longitudinal, Mendelian randomization

## Abstract

**Background:** DNA methylation is associated with body mass index (BMI), but it is not clear if methylation scores are biomarkers for extant BMI, or predictive of future BMI. Here we explore the causal nature and predictive utility of DNA methylation measured in peripheral blood with BMI and cardiometabolic traits.

**Methods:** Analyses were conducted across the life course using the ARIES cohort of mothers (n=792) and children (n=906), for whom DNA methylation and genetic profiles and BMI at multiple time points (3 in children at birth, in childhood and in adolescence, 2 in mothers during pregnancy and in middle age) were available. Genetic and DNA methylation scores for BMI were derived using published associations between BMI and DNA methylation and genotype. Causal relationships between methylation and BMI were assessed using Mendelian randomisation and cross-lagged models.

**Results:** The DNA methylation scores in adult women explained 10% of extant BMI variance. However, less extant variance was explained by scores generated in the same women during pregnancy (2% BMI variance) and in older children (15-17 years; 3% BMI variance). Similarly, little extant variance was explained in younger children (at birth and at 7 years; 1% and 2%, respectively). These associations remained following adjustment for smoking exposure and education levels. The DNA methylation score was found to be a poor predictor of future BMI using linear and cross-lagged models, suggesting that DNA methylation variation does not cause later variation in BMI. However, there was some evidence to suggest that BMI is predictive of later DNA methylation. Mendelian randomisation analyses also support this direction of effect, although evidence is weak. Finally, we find that DNA methylation scores for BMI are associated with extant cardiometabolic traits independently of BMI and genetic score.

**Conclusion:** The age-specific nature of DNA methylation associations with BMI, lack of causal relationship, and limited predictive ability of future BMI, indicate that DNA methylation is likely influenced by BMI and might more accurately be considered a biomarker of BMI and related outcomes than a predictor. Future epigenome-wide association studies may benefit from further examining associations between early DNA methylation and later health outcomes.

## Background

Obesity has a considerable burden on healthcare and has been shown to be predictive of mortality [1]. In recent years there has been a gradual increase in body mass index (BMI) in many countries [2] and interventions to decrease BMI have had limited success [3,4]. Predicting BMI early on and performing targeted interventions is an alternative strategy that could be more effective.

The aetiology of BMI comprises both genetic and environmental factors, with heritability likely below 0.5 [5,6]. It has been suggested that natural variation in DNA methylation levels may be a risk factor for certain diseases and play a role in the phenotypic variation of many traits [7–9]. In some cases, DNA methylation may provide the molecular link between environmental factors and associated disease risk, for example, recent studies have suggested that DNA methylation may be the mechanism allowing environmental factors or increased BMI to lead to obesity-related health outcomes [10–12]. Therefore, it could be useful as a predictor of such health outcomes.

Recent studies [13–15] suggest that DNA methylation associates with BMI trait variance independent of genetic variation. Associated genes have been shown to be involved in processes such as metabolism, inflammation, metabolic disease and cardiovascular disease amongst others. This suggests that associated DNA methylation loci may belong to causal pathways linking BMI and metabolic, cardiovascular and other obesity-related health outcomes, but this requires further exploration.

Since genetic variants are fixed at conception, genetic variants associated with BMI can be used early in life to predict later BMI [16]. The relationship of DNA methylation at BMI associated loci, however, is more complex. Methylation levels vary over time and may change in response to environmental or phenotypic changes so earlier methylation variation is not guaranteed to predict later BMI levels. Recent work [14] has suggested that change in BMI is more likely to be causal for changes in DNA methylation than vice versa. This would suggest that current DNA methylation scores are simply *biomarkers* for extant BMI. However, the utility of DNA methylation as a *predictor* for future trajectories of BMI would be of considerably greater utility.

The first aim of our study was to investigate if there is a temporal association between DNA methylation and BMI. We approached this question by using genome-wide DNA methylation profiles from the Accessible Resource for Integrated Epigenomics Studies (ARIES) [17] subset of the Avon Longitudinal Study of Parents and Children (ALSPAC) [18,19]. ALSPAC is a prospective cohort of children born in the former county of Avon, England during 1991 and1992. DNA methylation profiles were generated in children from blood collected at three time points (birth, childhood, adolescence) and from their mothers at two time points (during pregnancy and at middle-age). We used these DNA methylation profiles along with multiple measurements of BMI genetic profiles to determine if DNA methylation predicted BMI later in life, independently of genetic variation and BMI itself, and vice versa. In doing so, our objective was to determine if DNA methylation is a predictor for BMI or simply a biomarker that proxies current or previous BMI values.

DNA methylation scores for BMI may be useful for the detection of adverse health outcomes related to BMI. For example, *Wahl et al* found that DNA methylation could be used to identify individuals at high risk of incident type 2 diabetes, independently of other explanatory factors, including BMI itself [14]. Therefore, the second aim of our study was to see if DNA methylation scores for BMI contributed anything above BMI itself in predicting related adverse health outcomes.

## Methods

### Cohort description

We used children and mothers data from the Avon Longitudinal Study of Parents and Children (ALSPAC) cohort in this study [18,19]. The ALSPAC cohort is a prospective birth cohort study in which 14,541 pregnant women living in Avon, UK, with an expected delivery date from 1st April 1991 to the 31st December 1992 were initially recruited. Of these, 13,988 children were still alive 1 year later and have been followed-up with regular questionnaires and clinical measures, providing behavioural, lifestyle and biological data. When the children were approximately 7 years of age, an attempt was made to bolster the initial sample with eligible cases who had failed to join the study originally. As a result, when considering variables collected after age 7, the total sample size for those alive at 1 year of age is 14,901. The study website contains details of all the data that is available through a fully searchable data dictionary and variable search tool http://www.bristol.ac.uk/alspac/researchers/our-data/.

We only included participants that were also in the sub-study Accessible Resource for Integrated Epigenomic Studies (ARIES), where methylation data was available for these individuals [17]. After excluding those without methylation or phenotypic data and those that had withdrawn consent, we had data available for analyses from 823 children at birth, 906 for childhood (age 7), 770 for adolescence (age 15) and 792 for pregnant mothers and 726 for middle-aged mothers. The mean age and BMI are presented in **Table 1**. Sex is also included for children only, as adults were all female.

**Table 1.** Cohort description

|  | **Children** |  |  | **Mothers** |  |
| --- | --- | --- | --- | --- | --- |
|  | **Birth**<br>**(n=823)** | **Childhood**<br>**(n=906)** | **Adolescence**<br>**(n=770)** | **Pregnancy**<br>**(n=792)** | **Middle-age**<br>**(n=726)** |
| **Mean age in years (SD)** | NA | 7.45 (0.15) | 17.35 (0.88) | 28.87<br>(4.30) | 47.66<br>(4.26) |
| **Percentage of females** | 51.03% | 50.44% | 53.38% | 100% | 100% |
| **Mean BMI (kg/m<sup>2</sup>) or birthweight (grams) (SD)** | 3490.93<br>(477.76) | 16.20<br>(2.04) | 22.58 (3.83) | 22.72<br>(3.63) | 26.41<br>(5.09) |

*SD = Standard Deviation, BMI= Body Mass Index*

### Ethics

Ethical approval for the study was obtained from the ALSPAC Ethics and Law Committee and the Local Research Ethics Committees. Informed consent for the use of data collected via questionnaires and clinics was obtained from participants following the recommendations of the ALSPAC Ethics and Law Committee at the time.

### Phenotypic measures

The time points investigated in this study are pregnancy and middle-age data for the mothers and birth, childhood (age 7) and adolescence (age 15-17 for methylation data and age 17 for BMI data) data for the children.

Measures of height and weight were collected at research clinic visits. Height was measured to the nearest millimetre using the Harpenden Stadiometer. Weight was measured using the Tanita Body Fat Analyser to the nearest 50g. BMI was calculated by dividing weight (kilograms) by height (meters) squared (kg/m^2^). For measurements at birth, birth weight was collected and is used here instead of BMI.

Data for smoking and highest education level of mothers, age, sex, sample type and cell type proportions (B cell, CD4T, CD8T, Gran, Mono and NK) were used as covariates in analyses (for children, information on matched mothers smoking and maternal education were used). Various answers from questions regarding smoking in mothers (in pregnancy and currently) were used, from questionnaire data. These included the number of cigarettes smoked per day before pregnancy, during the first three months of pregnancy and the number of cigarettes smoked in the last 2 weeks during or just after pregnancy. These measures were combined to create a variable for whether the mother smoked in pregnancy. We also used data on the number of cigarettes smoked per day, whether they responded as being a current smoker and the time passed since they stopped smoking if this was within the last 12 months to create a variable for current or recent (within the last 12 months) smoker. Education level for the mother’s was also taken from questionnaire data, where participants were asked “What educational qualifications do you, your partner, your mother, and your father have?”. They were asked to select all options that applied to them and we used the highest education qualification for the participant. The options for this were CSE/none, vocational, O level, A level or degree.

Variables for cardiovascular outcomes for middle-aged mothers and adolescents were also used. These variables were from blood samples, obtained during clinic visits and included fasting glucose and insulin, triglycerides and low-density lipoprotein (LDL). Sitting diastolic and systolic blood pressures (SBP) from the right arm were collected during clinic visits. An Omron M6 upper arm blood pressure/pulse monitor was used to take 2 readings of blood pressure and then mean values were used. The data for triglycerides, glucose and insulin were skewed so we log transformed this data for use in analyses.

### Methylation data

Methylation profiling in the ARIES subset was conducted using DNA samples from blood taken at clinic visits or after delivery from the umbilical cord in the case of the birth time point. Blood from 1,018 mother–child pairs (children at three time points and their mothers at two time points) were selected for analysis as part of the Accessible Resource for Integrative Epigenomic Studies (ARIES, http://www.ariesepigenomics.org.uk/) [17]. Following DNA extraction, samples were bisulphite converted using the Zymo EZ DNA Methylation™ kit (Zymo, Irvine, CA, USA). Following conversion, genome-wide methylation was measured using the Illumina Infinium HumanMethylation450 (HM450) BeadChip. The arrays were scanned using an Illumina iScan, with initial quality review using GenomeStudio. ARIES data were pre-processed and normalised using the *meffil* R package [20]. ARIES consists of 5,469 DNA methylation profiles obtained from 1,022 mother-child pairs measured at five time points. Low quality profiles were removed from further processing, and the remaining 4,593 profiles were normalised using the Functional Normalization algorithm [21] with the top 10 control probe principal components. From the ARIES dataset, sample type and normalised methylation data was extracted and cell type proportion data were estimated using the Houseman method [22]. Full details of the pre-processing and normalization of ARIES has been described previously [20].

### Genotyping

Genetic data were collected from blood samples obtained in clinic visits. Genotyping was conducted with the Illumina HumanHap550 quad chip for children and the Illumina human660W-quad array for mothers. Quality control measures were carried out and haplotypes estimated using ShapeIT. A phased version of the 1000 genomes reference panel from the Impute2 reference data repository was used and Imputation of the target data was performed with this, using all reference haplotypes. A large proportion of the cohort has genome-wide data from these samples and a subset of this data is used in this study [18].

### Genetic and epigenetic scores

To investigate whether reported DNA methylation associations with BMI could be observed in an independent cohort, we calculated DNA methylation scores from published effect sizes for 135 CpG sites from the *Mendelson et al* [15] meta-analysis of DNA methylation and BMI. Scores were obtained for each ARIES methylation profile by multiplying the CpG site methylation levels in that profile with the corresponding published effects estimates and then summing the products.

Genetic scores were similarly derived using effect sizes for 97 SNPs from the GIANT consortium BMI genome-wide association study (GWAS) [23]. Scores were created using Plink V1.9 (https://www.cog-genomics.org/plink2) with the score and sum commands, however one of the SNPs did not meet imputation score filters (rs12016871), so the score was constructed using only 96 SNPs.

To perform simplified versions of Mendelian randomisation (MR), we used summary statistics from methylation quantitative trait loci (mQTL). mQTLs are genetic variants associated with DNA methylation [24]. To identify mQTLs we looked these up in mQTLdb [25], which contains mQTLs below a conservative p-value threshold of 1e-07. If multiple mQTLs were identified for an individual CpG site, the one with the lowest p-value reported in the GIANT study for BMI was used as the mQTL for the MR analysis. If these mQTLs were unavailable, then proxies of these SNPs were obtained. These were SNPs with the next lowest p-value for that CpG, which were also present in the BMI GWAS data. We used the last p-value available in the BMI GWAS for each mQTL to maximise power. Of the 135 CpG sites we queried, 89 had an instrument at this threshold.

## Statistical analysis

### Observational associations at the same time point

Linear regression models, with adjustments for covariates, were used to test observational associations. When testing for association between genetic and/or methylation scores of BMI, BMI was the dependent variable and the methylation and/or genetic score the independent variables. For models including a genetic score, age was included as a covariate. For models including a methylation score, the covariates included were age, sample type and estimated blood cell type proportions. Sex was additionally included as a covariate in all models analysing child data. For predicting BMI, BMI was used as the dependent variable, and when predicting methylation, methylation score was used as the dependent variable.

To compare the relative contributions of genetic score and methylation score to BMI, an analysis of variance (ANOVA) test was carried out comparing the following three models, with the full model (model 3) being compared to each of the reduced models (models 1 and 2):

1. BMI ∼ methylation score + covariates
2. BMI ∼ genetic score + covariates
3. BMI ∼ methylation score + genetic score + covariates

Finally, we investigated how BMI and DNA methylation change over the life course. Firstly, we calculated correlations of BMI and DNA methylation score at different time points for mothers and children separately. We then examined the correlation of BMI and DNA methylation scores between paired children and mothers at the different time points. Thirdly, we also calculated correlations for all individual CpG sites across the different timepoints and between paired child’s cord blood DNA and mother’s antenatal peripheral blood DNA values.

### Observational associations across the life course

To investigate whether DNA methylation might be predictive of later BMI or vice versa, we assessed associations between different time points in mothers and children, using linear regression models, similar to those used in the observational analyses within the same time point.

### Exploration of a causal relationship between BMI and DNA methylation

#### Cross-lagged model

We analysed the temporal relationship of BMI and DNA methylation, using a cross-lagged model. This approach allows exploration of the relationships between earlier BMI and later methylation score in two separate systems, one in the children (using childhood and adolescence) and one in the mothers (using the antenatal and middle age time points). The R package OpenMx (version 2.13.2) [26] was used to build a cross-lagged model, shown in Figures 3 and 4. Values for each of the free parameters or paths are estimated in the model. The paths were from earlier BMI to methylation at the same time point, later methylation and later BMI; and from earlier methylation to later BMI and methylation; and from later BMI to methylation at the same time point. Each path was sequentially tested in a sub model analysis, where that path was fixed to 0 and this sub model was compared against the full model using a likelihood ratio test. If a sub model had a significantly worse fit then that path was retained, but otherwise dropped because it was not important to the overall system.

**Figure 1.**
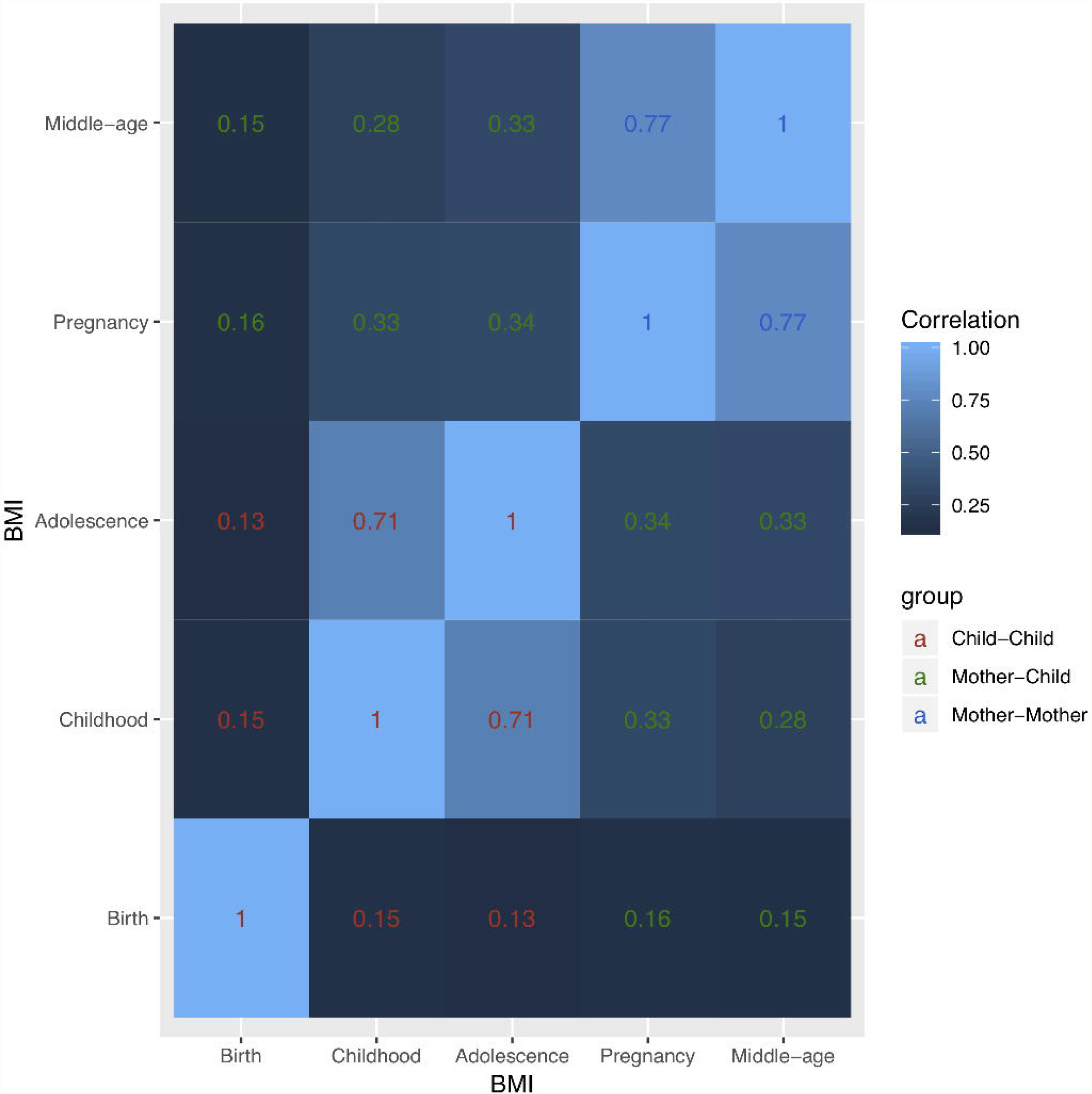
Correlation matrix of BMI in children (birth, childhood, adolescence) and mothers (pregnancy and middle-age). Legend: This correlation matrix shows the correlations over time for BMI at all time points in children and mothers. There is a correlation of BMI in children and mothers over time and between paired mother and children’s BMI.

**Figure 2.**
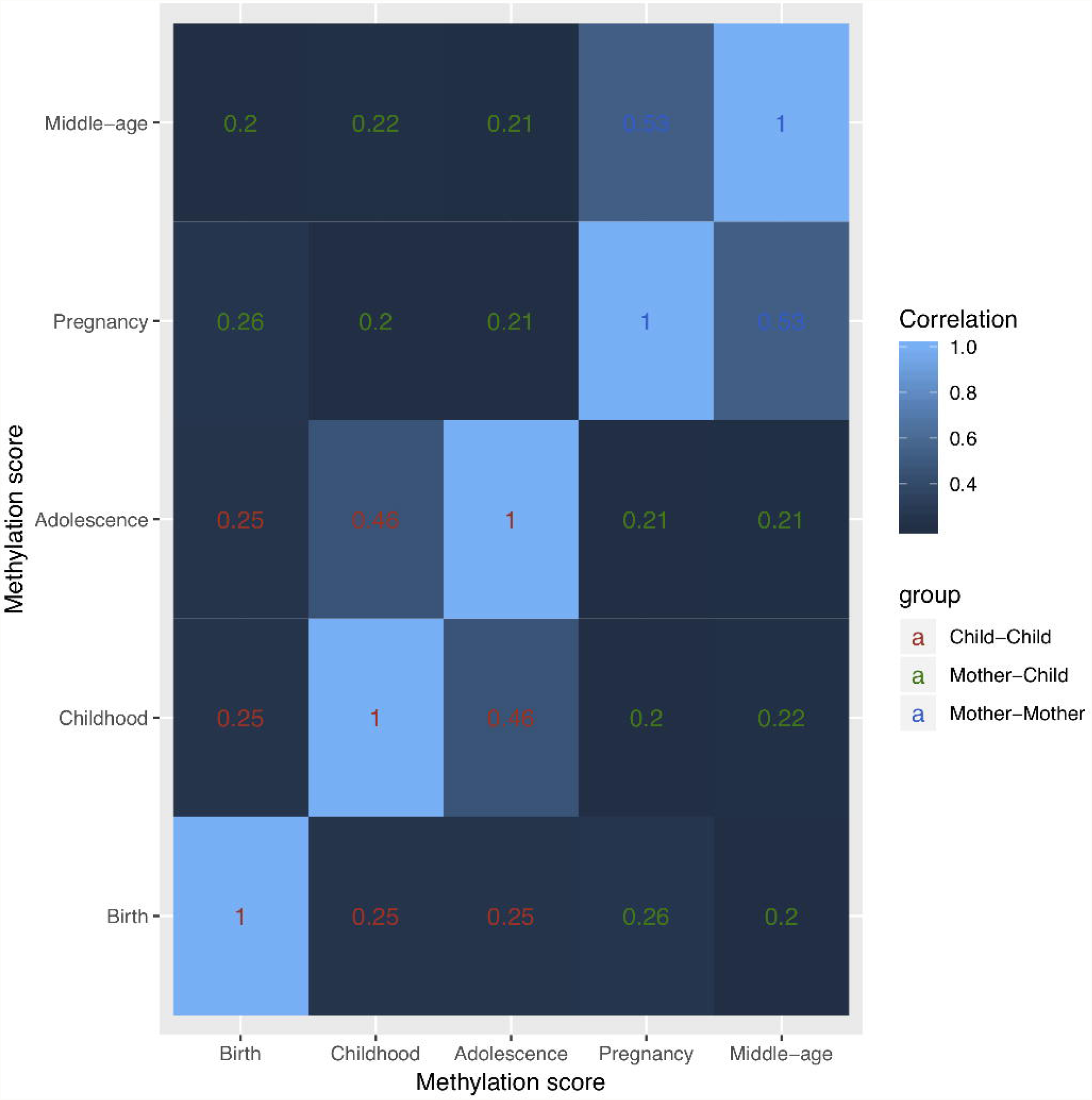
Correlation matrix of methylation score in children (birth, childhood, adolescence) and mothers (pregnancy and middle-age). Legend: This correlation matrix shows the correlations over time for overall methylation score at all time points in children and mothers. There is a correlation of methylation score in children and mothers over time and between paired mother and children’s methylation scores.

**Figure 3.**
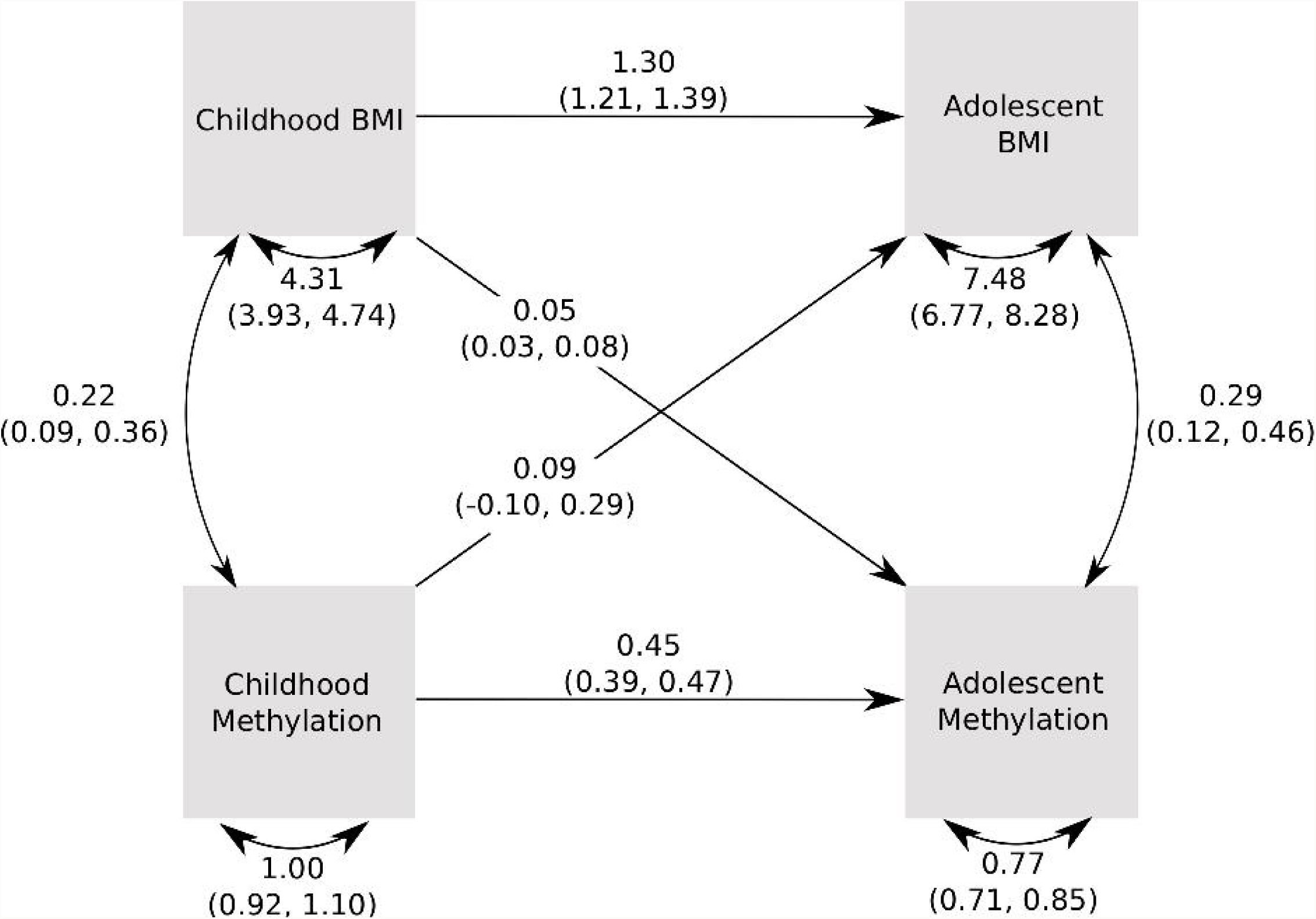
Pathway diagram for the cross-lagged model for childhood and adolescence. Legend: This diagram shows the observed variables in boxes. Single headed arrows indicate linear regressions and double headed, curved arrows indicate variances/covariances. Estimates for the linear relationships are shown on the arrows, as are the values for variances and covariances.

**Figure 4.**
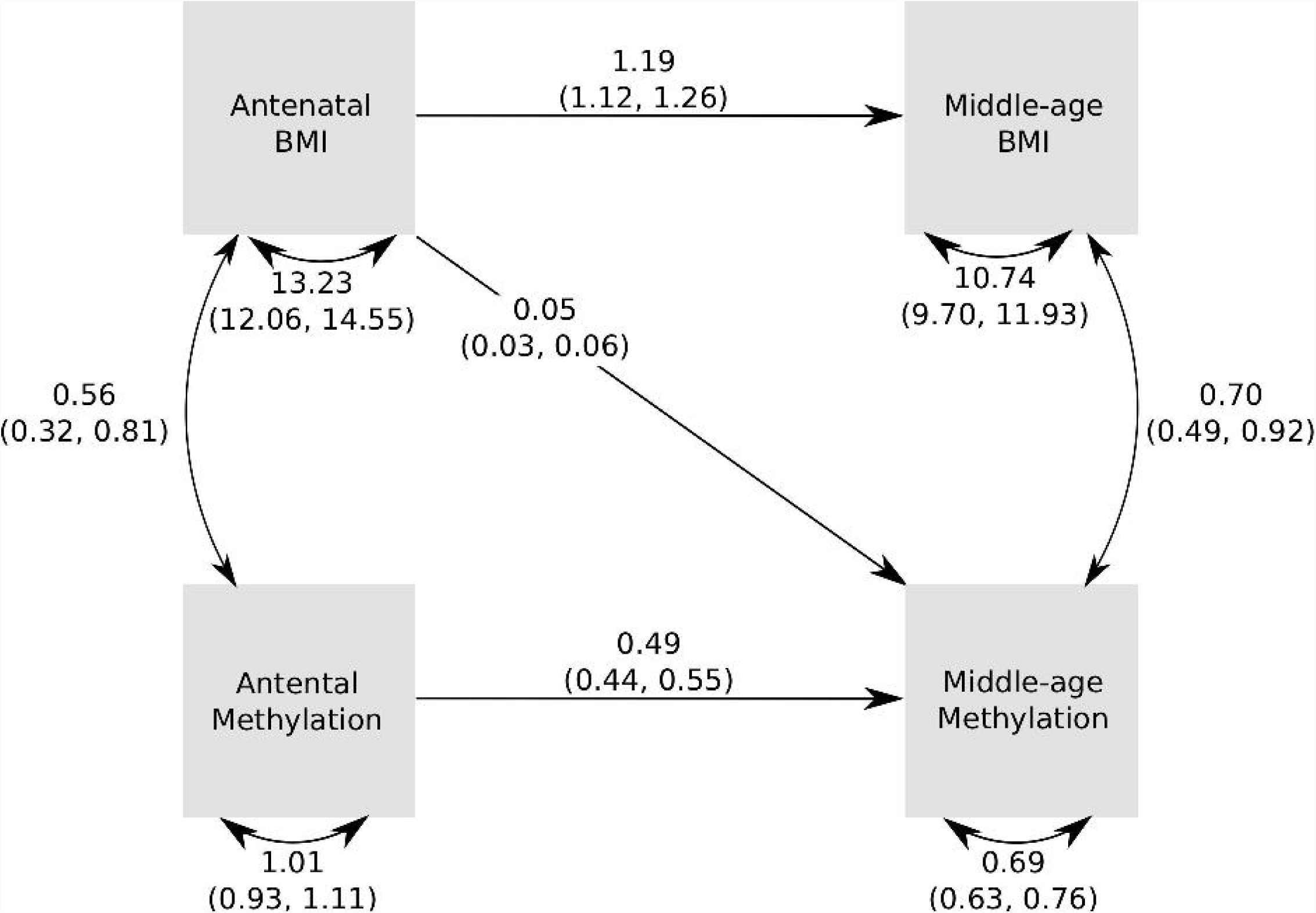
Pathway diagram for the cross-lagged model for pregnancy and middle-aged mothers. Legend: This diagram shows the observed variables in boxes. Single headed arrows indicate linear regressions and double headed, curved arrows indicate variances/covariances. Estimates for the linear relationships are shown on the arrows, as are the values for variances and covariances.

#### Mendelian randomisation

To investigate causal inference more directly, an approach based on MR was adopted. To test if changes in BMI cause changes in DNA methylation, we calculated genetic scores for BMI, as previously described, and tested the association of this score with each of the 135 BMI-associated CpG sites. A Fisher’s test was then applied to combine the association p-values for all 135 association tests. To increase power, the adolescent and middle-aged mother’s data was subsequently combined, and the association was tested again using a mixed model to account for relatedness.

Two-sample MR was applied to explore the reverse direction, i.e. a causal effect of DNA methylation on BMI. In this approach, summary statistics from the BMI GWAS were obtained for the mQTLs (or proxies of these, if these SNPs were unavailable) for the 135 BMI-associated CpG sites.

### Confounder analysis

To investigate whether any associations found between methylation and BMI were due to confounding by smoking or education, we compared linear models of BMI and DNA methylation with and without smoking (prenatal smoking during pregnancy for children and own smoking for adults) and education as covariates.

### Cardiovascular trait analyses

Linear regression models were used to test observational associations between the methylation and genetic scores for BMI against cardiovascular outcomes. These were performed with and without adjustment for BMI and using other covariates, as with other models. ANOVA tests were used to compare the relative contributions of BMI and the genetic and methylation scores and results are reported for these comparisons.

## Results

### Establishing associations between DNA methylation and BMI within time-points

The methylation score for BMI, derived from the *Mendelson et al* epigenome-wide association study (EWAS) [15], was strongly associated with BMI in middle aged mothers, explaining 10% of the variation in BMI (p=1.58E-23). The association was weaker in mothers during pregnancy (2% variance explained, p=2.31E-06) and in children at birth (2% variance explained, p=6.83E-05), childhood (1% variance explained, p=2.23E-04) and adolescence (3% variance explained, p=3.91E-11). Full results are presented in **Table 2**.

**Table 2.** Associations between methylation and genetic scores and BMI/birth weight at the same time point.

|  | N | Associations between methylation score and BMI/birth weight |  |  | Associations between genetic score and BMI/birth weight |  |  |
| --- | --- | --- | --- | --- | --- | --- | --- |
|  |  | Effect size<sup>1</sup>, in kg/m<sup>2</sup> per SD in methylation score (CI) | p-value | Adjusted R-squared<sup>3</sup> | Effect size<sup>2</sup> (CI) | p-value | Adjusted R-squared<sup>3</sup> |
| **Birth** | 823 | 69.27 (35.35, 103.18) | 6.83E-05 | 0.02 | -219.75 (-412.59, -26.90) | 0.03 | 0.004 |
| **Childhood** | 906 | 0.25 (0.12, 0.39) | 2.23E-04 | 0.01 | 2.24 (1.45, 3.02) | 2.72E-08 | 0.03 |
| **Adolescence** | 770 | 0.97 (0.69, 1.26) | 3.91E-11 | 0.03 | 5.25 (3.65, 6.85) | 2.12E-10 | 0.05 |
| **Pregnancy** | 792 | 0.62 (0.36, 0.87) | 2.31E-06 | 0.02 | 2.91 (1.38, 4.43) | 1.95E-04 | 0.02 |
| **Middle-age** | 726 | 2.06 (1.67, 2.46) | 1.58E-23 | 0.10 | 4.72 (2.54, 6.90) | 2.45E-05 | 0.02 |

<sup>1</sup> From model adjusting for age (except at birth), sex (where applicable), sample type
(where applicable), and cell type proportions (B cell, CD4T, CD8T, Gran, Mono and NK).
<sup>2</sup> From model adjusting for age and sex (where applicable)
<sup>3</sup> Adjusted R-squared obtained from model with only the methylation score or genetic score
and no other covariates.

Genetic scores for BMI, derived from published GWAS summary statistics [23], were also associated with BMI, with the strongest association found for children in adolescence with 5% variance explained in BMI (p=2.12E-10) and weaker associations found at all other time points (see **Table 2** for full results).

The genetic and methylation score associations appear to be mostly independent (**Table 3**), as the combined model with both genetic and methylation scores performed better than both the methylation score (ANOVA test p-values ranged from 1.93E-10 to 0.04) and genetic score models (ANOVA test p-values ranged from 4.55E-21 to 1.28E-03) alone for all time points. This validates previous findings that the genotype and DNA methylation explain independent subsets of BMI variation [13,15].

**Table 3.** Results from combined model and ANOVA comparing this with models for methylation and genetic scores.

|  | Combined model<br>adjusted R-squared<sup>1</sup> | ANOVA (model<br>1 vs model 3) | ANOVA (model 2<br>vs model 3) |
| --- | --- | --- | --- |
| **Birth** | 0.03 | 0.04 | 1.56E-09 |
| **Childhood** | 0.04 | 9.03E-09 | 1.28E-03 |
| **Adolescence** | 0.08 | 1.93E-10 | 5.28E-11 |
| **Pregnancy** | 0.04 | 2.22E-04 | 5.36E-05 |
| **Middle-age** | 0.12 | 2.77E-05 | 4.55E-21 |

<sup>1</sup> Adjusted R-squared obtained from model with both the methylation and the genetic scores and no other covariates
Model 1 includes the methylation score, model 2 includes the genetic score and model 3 is the combined model including the methylation and genetic score. All models also included the relevant covariates

### Stability of phenotypic values over time and between generations

We evaluated the extent to which individual BMI levels correlated over time and between mothers and children. The strongest BMI correlations were observed in children between age 7 and adolescence and in mothers between pregnancy and middle age (R ∼ 0.7). Intermediate correlations were observed between children and mothers at all time points (R ∼ 0.3) except birth. Lowest BMI correlations (R ∼ 0.15) were observed with birth, likely because birthweight is a different measure than BMI (**Figure 1**).

The BMI methylation score correlations exhibited similar patterns but were generally lower than for BMI. Strongest correlations were observed in children between age 7 and adolescence and in mothers between pregnancy and middle age (R ∼ 0.5). All other correlations were between 0.2 and 0.25. Thus, methylation scores at birth were more highly correlated with later time points and with maternal methylation scores than birthweight and BMI (**Figure 2**). Given the weak association of methylation in childhood with BMI, factors other than BMI likely contribute to the correlation of DNA methylation over time and between mothers and children.

Finally, to examine whether there are particular CpG sites that correlate more strongly over time and between paired children and mothers, we tested the correlation of each site at different time points (Supplementary Table 1, Additional file 1). We observe that median correlations across all CpG sites follow a similar pattern to correlations of the methylation scores over time, where the strongest correlations were observed in children between age 7 and adolescence and in mothers between pregnancy and middle age (R ∼ 0.2). Only these two sets of timepoints have CpG sites with correlation R > 0.5. There are six such CpG sites for each and two of these are common to both (cg16611584, cg24145109). Both of these CpG sites are highly correlated across all time points.

### Predicting future BMI with past DNA methylation scores

We next investigated whether the BMI methylation score was predictive of BMI at later timepoints (**Table 4**) or vice versa (if BMI was predictive of methylation score at later timepoints; **Table 5**). We observed some evidence that methylation score in childhood could be predictive of BMI in adolescence (p=0.004), although the association disappeared when adjusting for childhood BMI (p=0.20) and there was stronger evidence for the converse, that is BMI in childhood predicting adolescent DNA methylation (p=1.52E-06 even when adjusting for childhood methylation score). We observed the same in the mothers between pregnancy and middle age; that is, the association between antenatal (earlier) methylation score and middle-age (later) BMI (p=0.02) essentially disappears when adjusting for antenatal (earlier) BMI (p=0.13). Also, the association between antenatal (earlier) BMI and middle age (later) DNA methylation is much stronger even when adjusting for antenatal (earlier) methylation score (p=5.48E-11).

**Table 4.**
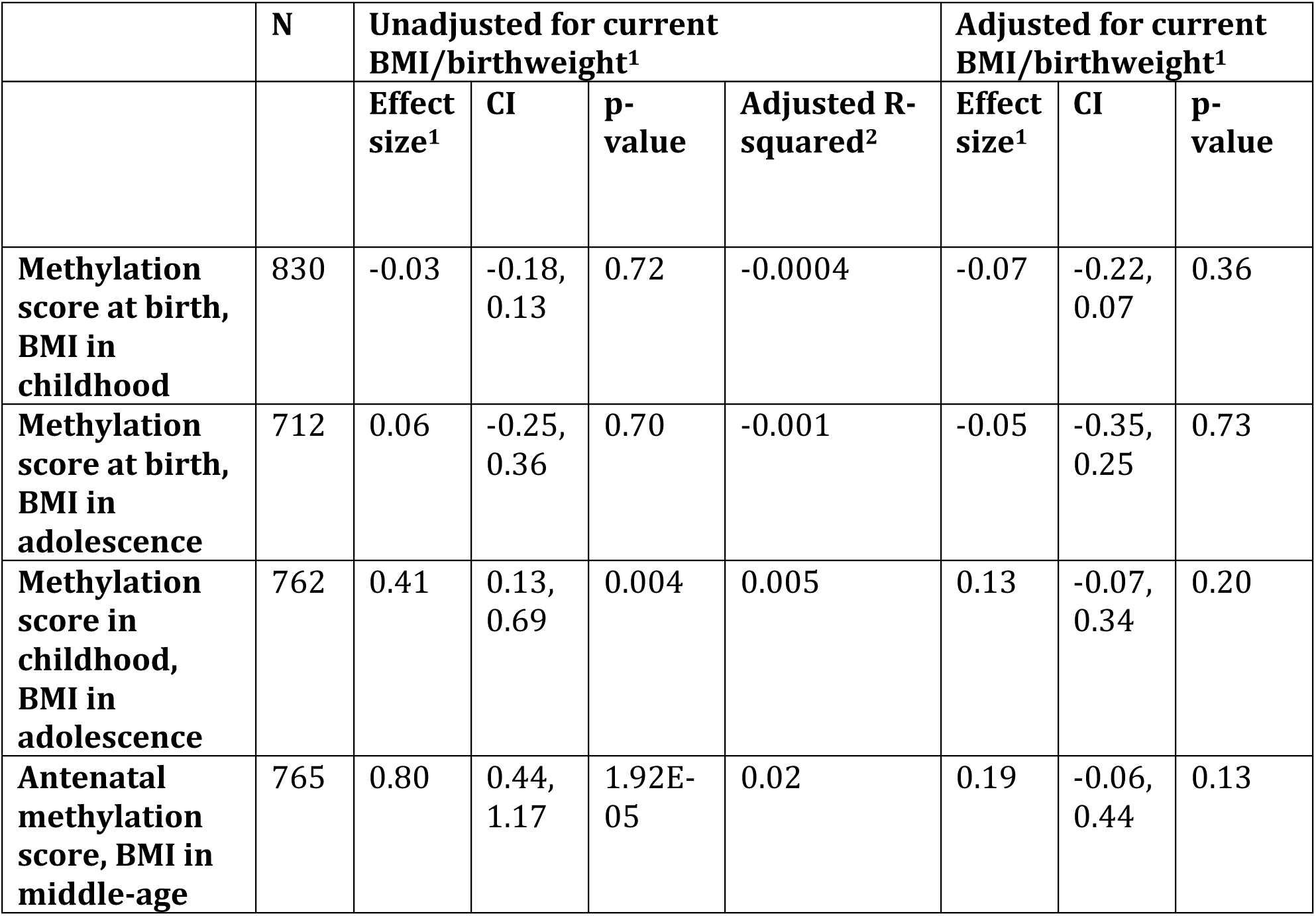
Associations between methylation score and BMI at later time points.

**Table 5.** Associations between BMI/birthweight and methylation score at later time points.

|  | N | Unadjusted for current methylation score<sup>1</sup> |  |  |  | Adjusted for current methylation score<sup>1</sup> |  |  |
| --- | --- | --- | --- | --- | --- | --- | --- | --- |
|  |  | Effect size<sup>1</sup> | CI | p-value | Adjusted R-squared<sup>2</sup> | Effect size<sup>1</sup> | CI | p-value |
| **Birthweight, methylation score in childhood** | 814 | -0.0001 | -0.0002, 0.00004 | 0.17 | 0.003 | -0.0002 | -0.0003, -0.00006 | 0.006 |
| **Birthweight, methylation** | 812 | 0.000003 | -0.0001, 0.0001 | 0.96 | 0.002 | -0.00008 | -0.0002, 0.00005 | 0.22 |
| **score in adolescence** |  |  |  |  |  |  |  |  |
| **BMI in childhood, methylation score in adolescence** | 891 | 0.09 | 0.06, 0.12 | 3.05E-09 | 0.02 | 0.06 | 0.04, 0.09 | 1.52E-06 |
| **Antenatal BMI, methylation score in middle-age** | 851 | 0.07 | 0.05, 0.08 | 1.42E-16 | 0.06 | 0.05 | 0.03, 0.05 | 5.48E-11 |

<sup>1</sup> From model adjusting for age (except at birth), sex (where applicable), sample type
(where applicable), and cell type proportions (B cell, CD4T, CD8T, Gran, Mono and NK).
<sup>2</sup> Adjusted R-squared obtained from model with only the methylation score and no other
covariates

### Exploration of temporal relationships between BMI and DNA methylation

To further evaluate the temporal associations between DNA methylation and BMI, we used cross-lagged models to test which paths from earlier trait measures and scores were important for later trait measures and scores. Agreeing with the results from adjusted linear models, these did reveal a pathway between BMI in childhood and methylation score in adolescence. **Figure 3** shows the estimates and variances/covariances obtained from the main model. The only path that could be dropped from the model without affecting model fit was between childhood methylation and adolescent BMI (p=0.35 for this path and p < 1.79E-4 for all other paths; Supplementary Figure 1, Additional file 1). Cross-lagged model fits in mothers (**Figure 4**) also revealed a pathway from (earlier) BMI in pregnancy to (later) DNA methylation in middle-age. The only path that could be dropped from the model without affecting model fit was from DNA methylation in pregnancy (earlier) and BMI in middle-age (later) (p=0.20 for this path and p<3.65E-09 for all other paths; Supplementary Figure 2, Additional file 1).

### Mendelian randomisation does not support a causal relationship of DNAm on BMI

We used two-sample MR to explore causal relationships between DNA methylation and BMI (Supplementary Table 2, Additional file 1). Using the BMI genetic score as an instrumental variable for BMI, we found little evidence for a causal link of BMI on each of the 135 CpG sites used to construct the methylation score (p-value range: 1.63E-03 – 9.99E-01, for all timepoints, with a Bonferroni-adjusted p-value threshold of 3.7E-04). A combined p-value for all 135 CpG sites obtained using Fisher’s method similarly indicated no strong association between the genetic score and methylation (p-value range: 0.82 – 1.00, for all timepoints). Furthermore, even when this analysis was repeated with a mixed model including data from both adolescents and middle-aged mothers to increase the power (Supplementary Table 3, Additional file 1), there was still little evidence of association (p-value range: 2.13E-02 – 9.98E-01).

The reverse causal direction, methylation variation causing BMI variation, was investigated using mQTLs for the 135 methylation score CpG sites as instrumental variables. These individual tests did not indicate a causal link (Bonferroni-adjusted p-value threshold of 3.7E-04; Supplementary Table 4, Additional file 1) although combining the test p-values using Fisher’s method did provide weak evidence for a causal association (Fisher’s p-value = 0.03).

### Confounder analyses

Sensitivity analyses showed that associations between BMI and methylation score remained unaffected by the inclusion of potential confounders in the majority of models (Supplementary Table 5, Additional file 1). Smoking and education appeared to be associated with methylation score in some models, however, most of the associations between BMI and methylation score survived these adjustments (p < 0.007 = 0.05/7).

### Associations with cardiovascular traits

Finally, given that BMI is a risk factor for cardiovascular disease, we tested within-timepoint associations between the DNA methylation and genetic scores for BMI and cardiovascular traits to examine if the specificity of these scores. Firstly, we observed that methylation associations were partially independent of BMI for LDL (p=0.02) and glucose (p=0.03) in adolescence and triglycerides (p=3.00E-03), LDL (p=0.01) and SBP (p=0.05) in mothers at middle age. Similarly, we observed that methylation associations were partially independent of genetic effects on SBP in adolescence (p=0.05) and triglycerides in mothers (p=3.27E-03) (Supplementary Table 6, Additional file 1). We also observed that genetic effects were partially independent of BMI for LDL (p=0.03), glucose (0.05) and SBP (0.03) in adolescence and triglycerides (2.01E-04), LDL (0.02) and SBP (0.02) in mothers. Finally, we observed that the genetic effects were partially independent of methylation and BMI for LDL (p=0.02) and glucose (p=0.03) in adolescence and triglycerides (p=3.80E-03), LDL (p=0.01) and SBP (p=0.02) in mothers (Supplementary Table 7, Additional file 1).

## Discussion

In this study we have demonstrated strong associations between DNA methylation and genetic scores for BMI [15] in both adults and children. Importantly, the use of temporal data indicates that the DNA methylation scores are not predictive. While the association between DNA methylation scores and BMI within time point are strong, the associations between earlier methylation scores later BMI are weak, and these signals do not improve the model of simply using earlier BMI as a predictor for later BMI. Hence, it may be more appropriate to term the DNA methylation score as a biomarker rather than a cause or predictor of BMI.

We observed within-timepoint associations between DNA methylation score for BMI and health outcomes for which BMI is a risk factor. These associations were independent of BMI and genotype, suggesting that the DNA methylation – BMI associations might arise due to unmeasured confounders that also influence those outcomes. DNA methylation could be used as a biomarker for these outcomes, above and beyond BMI and genetic variation.

Our study builds upon previous research in this field showing that both genetic and environmental factors contribute to variance in BMI [5,23]. Previous work investigating the relationship between DNA methylation and BMI has found associations with specific DNA methylation sites [10–12]. However results from these studies are fairly inconsistent and practical implications of these associations have not yet been identified [27]. In extension of this previous research we have used 135 CpG sites, identified in an EWAS meta-analysis [15], to create methylation scores that are also associated with BMI in an independent cohort. The main novel component of our study is that we look across the life course in children and adults. Another study investigating DNA methylation profile and obesity in children aged 6-14 years also found an association between childhood obesity and a separate set of differentially methylated CpG sites, supporting our finding that DNA methylation is associated with BMI in adolescents [28]. Our findings in younger children were however weaker and could be due to basing our analysis on methylation at CpG sites associated with BMI in adults.

We have also investigated the nature of these associations further and found that there may be some predictive capability of earlier BMI to later DNA methylation at multiple time points, however we found no evidence to suggest this association is causal, which could be due to several reasons. Firstly, it may be that there is a lack of power in these analyses for MR to detect a causal relationship. The *Wahl et al* paper [14] suggests that with larger sample sizes the direction of effect is likely to be from BMI to DNA methylation. Therefore, it is possible that with a larger sample size we could confirm this direction of causality. It could also be that there are other unknown confounders, which are mediating this effect and future research should focus on investigating this further. Our MR analyses suggest a weak aggregate causal association from DNA methylation to BMI, however due to the DNAm instruments being enriched for genic regions, this association is unlikely to be stronger than expected against a more appropriate null that reflects that BMI is more strongly influenced by genetic variants in genic regions [23]. We also found associations of the BMI DNAm scores with cardiovascular outcomes. If BMI causes changes in DNAm, as our analyses seem to suggest, then DNAm changes may fall on the causal pathway between BMI and these cardiovascular outcomes. Therefore, DNAm scores may have potential use as predictors of cardiovascular outcomes, although generalisability to other populations would need to be confirmed. This has been suggested in previous studies for diabetes [14,29] and insulin-resistance [30].

### Limitations

This study is subject to a number of limitations. Firstly, whilst the size of the ARIES cohort is larger than or equivalent to samples used in other DNA methylation studies, it may still be too small to detect some associations, and this may be why we do not find any causal association for BMI to DNA methylation, as discussed above. The ARIES mQTL database used in the two-sample MR had a total sample size of 3,948, although this was split across three time points in approximately the same children and two time points in the mothers. We used the GIANT cohort for BMI in the two-sample MR and included 235,522 participants in these analyses. The GIANT sample size is much larger and therefore it is difficult to directly compare the strength of causal evidence in each direction between these samples. Secondly, whilst it is a strength of our study that we use multiple time points, in the mothers one of these is during pregnancy so these findings may not be generalizable outside of pregnancy, or generalizable to males. As with any epidemiological study there may also be measurement error present in our phenotypic data, for example smoking data was collected via self-report and there may be biases present in this data because of this. In addition, measurement error may be present in the genotyping data and methylation data. Finally, the DNA methylation data was collected from blood samples which may not be indicative of other tissue methylation levels such as adipose tissue which may be more relevant in a study looking at potential causal relationships between BMI and DNAm, however we assume the two may be correlated to an extent.

## Conclusions

In conclusion, our study finds that DNA methylation scores, as have so far been generated, are unlikely to be predictive of future BMI using earlier DNA methylation levels, and therefore is more appropriately considered a biomarker. This indicates that conducting EWAS using DNAm values and trait values measured at the same time point is not an effective strategy when attempting to create predictors. Therefore, future studies should perform EWAS that test for early DNAm values against later health outcomes to evaluate whether this may enable creating an effective predictor, which could then be tested in other populations.

## Abbreviations

BMI: Body mass index
ARIES: Accessible Resource for Integrated Epigenomic Studies
ALSPAC: Avon Longitudinal Study of Parents and Children
LDL: Low-density lipoprotein
SBP: Systolic blood pressure
MR: Mendelian randomisation
mQTL: methylation quantitative trait loci
ANOVA: Analysis of variance
EWAS: Epigenome-wide association study
GWAS: Genome-wide association study.

## Declarations

### Consent for publication

Not applicable

### Availability of data and material

The ALSPAC data management plan (http://www.bristol.ac.uk/alspac/researchers/data-access/documents/alspac-data-management-plan.pdf) describes in detail the policy regarding data sharing, which is through a system of managed open access. The steps below highlight how to apply for access to the data included in this paper and all other ALSPAC data. The datasets used in this analysis are linked to ALSPAC project number B2154; please quote this project number during your application.

- 1. Please read the ALSPAC access policy (PDF, 627 kB) which describes the process of accessing the data and samples in detail, and outlines the costs associated with doing so.
- 2. You may also find it useful to browse the fully searchable ALSPAC research proposals database, which lists all research projects that have been approved since April 2011.
- 3. Please submit your research proposal for consideration by the ALSPAC Executive Committee. You will receive a response within 10 working days to advise you whether your proposal has been approved.

If you have any questions about accessing data, please.

### Authors’ contributions

ZER, MJS, GH and CLR contributed to the planning and design of the analysis and drafting of this paper. OSPD advised on the use of the cross-lagged model design and contributed to final drafts of this paper.

## Acknowledgements

We are extremely grateful to all the families who took part in this study, the midwives for their help in recruiting them, and the whole ALSPAC team, which includes interviewers, computer and laboratory technicians, clerical workers, research scientists, volunteers, managers, receptionists and nurses.

## Additional files

Additional file 1: This contains supplementary tables S1-S7 and supplementary figures S1-S2. File format is .docx.

