## Additional file 1 for "The association of DNA methylation with body mass index: distinguishing between predictors and biomarkers"

**Supplementary table 1**. Table of correlations between all individual CpG sites across the different time points and between matched children (cord) and mothers (antenatal).

| **CpG** | **Weights** | **Correlations** | | | | |
| --- | --- | --- | --- | --- | --- | --- |
|  |  | **Cord and Age 7** | **Cord and Age 15** | **Age 7 and Age 15** | **Antenatal and Middle age** | **Children cord and Mothers antenatal** |
| **cg00108715** | 0.00733 | 0.045962819 | 0.018604982 | 0.069604459 | 0.153123118 | 0.007936365 |
| **cg00222799** | 0.00682 | 0.170421698 | 0.107590001 | 0.221024948 | 0.288764191 | 0.091416295 |
| **cg00574958** | -0.01773 | -0.017196773 | -0.016015899 | 0.104756763 | -0.005166276 | -0.008133243 |
| **cg01130991** | 0.01066 | 0.105781682 | 0.004965519 | 0.109968368 | 0.098710393 | 0.096059067 |
| **cg01243823** | -0.00805 | 0.169367175 | 0.16481023 | 0.519249018 | 0.4905285 | 0.021160999 |
| **cg01368219** | 0.00805 | 0.110430464 | 0.068545856 | 0.114992976 | 0.194533221 | 0.06293121 |
| **cg01455178** | 0.00927 | 0.11777125 | 0.081751744 | 0.284042742 | 0.214506147 | 0.08710877 |
| **cg01526748** | 0.00934 | 0.143456002 | 0.099076008 | 0.23264436 | 0.427982796 | 0.091762199 |
| **cg01597398** | -0.00595 | 0.267675998 | 0.206211267 | 0.23010843 | 0.240170713 | 0.047110777 |
| **cg01671681** | -0.00775 | 0.039850818 | 0.036226688 | 0.194923319 | 0.242617871 | 0.001668862 |
| **cg01751802** | 0.00815 | 0.066883599 | 0.13861652 | 0.261159291 | 0.337494336 | 0.012081673 |
| **cg01798813** | 0.01033 | 0.16944398 | 0.072445993 | 0.172455726 | 0.15328794 | 0.075248354 |
| **cg01881899** | 0.0112 | -0.004292765 | 0.000812628 | 0.066069917 | 0.049998625 | 0.021913283 |
| **cg02286155** | 0.00576 | 0.157029433 | 0.040307729 | 0.139734709 | 0.102256247 | 0.077612723 |
| **cg02571142** | 0.00837 | 0.155661334 | 0.193202619 | 0.33105896 | 0.361690873 | 0.076609151 |
| **cg02711608** | -0.00845 | 0.228940528 | 0.178863204 | 0.280630443 | 0.230067684 | 0.091502309 |
| **cg02716826** | -0.00762 | 0.088016848 | 0.103078513 | 0.194225409 | 0.196178098 | 0.08000903 |
| **cg03078551** | -0.01156 | 0.015470142 | 0.055231362 | 0.104600705 | 0.032101226 | 0.054821628 |
| **cg03218374** | 0.0085 | 0.050662191 | 0.050110145 | 0.254846374 | 0.250752662 | 0.051645351 |
| **cg03500056** | 0.00622 | 0.112632122 | 0.103587287 | 0.325716807 | 0.314633615 | 0.085776478 |
| **cg03682690** | 0.01002 | 0.093282075 | 0.055303315 | 0.23572367 | 0.261188632 | 0.047852226 |
| **cg03725309** | -0.00998 | 0.127407473 | 0.134195386 | 0.197995104 | 0.177136067 | 0.064769662 |
| **cg04011474** | -0.00815 | 0.045959681 | 0.076255522 | 0.053103648 | 0.23223476 | 0.013451447 |
| **cg04286697** | 0.00681 | 0.074478401 | 0.035824787 | 0.12980353 | 0.172390093 | 0.026169033 |
| **cg04483863** | 0.0087 | 0.086007948 | 0.072527511 | 0.054782533 | 0.113152964 | 0.05445323 |
| **cg04557677** | -0.00797 | 0.019866492 | 0.054264241 | 0.214092933 | 0.061449616 | -0.001837734 |
| **cg04816311** | 0.01005 | 0.080872804 | 0.121980496 | 0.184799835 | 0.25627928 | 0.070741029 |
| **cg04927537** | 0.01241 | 0.252808857 | 0.220315131 | 0.464633341 | 0.449798854 | 0.180616175 |
| **cg05119988** | -0.00925 | 0.04508268 | 0.114837356 | 0.156768972 | 0.121706432 | 0.048390872 |
| **cg06192883** | 0.008 | 0.081174855 | 0.098718807 | 0.150685797 | 0.201824782 | 0.024619121 |
| **cg06397161** | 0.01045 | 0.144622037 | 0.154786036 | 0.348486293 | 0.351147333 | 0.077138283 |
| **cg06500161** | 0.0164 | 0.101332931 | 0.057532419 | 0.196618803 | 0.14544405 | 0.105956013 |
| **cg06690548** | -0.0163 | 0.088518534 | 0.068929631 | 0.123909947 | 0.099679166 | 0.016021878 |
| **cg06876354** | 0.00649 | 0.077253641 | 0.134099997 | 0.168367095 | 0.194083082 | 0.164754393 |
| **cg06946797** | -0.00923 | 0.197799911 | 0.188765069 | 0.327269755 | 0.298997039 | 0.095915815 |
| **cg07012687** | 0.0078 | 0.08609032 | 0.163389255 | 0.279539515 | 0.211677893 | 0.117118741 |
| **cg07021906** | 0.0075 | 0.057380201 | 0.13496567 | 0.275349184 | 0.239289061 | 0.069490741 |
| **cg07037944** | -0.00733 | 0.238071862 | 0.225638332 | 0.270131601 | 0.184548769 | 0.087517482 |
| **cg07094298** | -0.01379 | 0.161549785 | 0.162044451 | 0.255792321 | 0.268992016 | 0.113893134 |
| **cg07504977** | 0.0102 | 0.122539086 | 0.106333944 | 0.217742383 | 0.146545287 | 0.086051738 |
| **cg07573872** | -0.00909 | 0.19884819 | 0.127491757 | 0.261554221 | 0.226732539 | 0.124157565 |
| **cg07730360** | 0.00722 | 0.146119731 | 0.052366811 | 0.091805383 | 0.076752935 | 0.042760448 |
| **cg07814318** | 0.00772 | 0.152441441 | 0.203805184 | 0.196608558 | 0.201739562 | 0.201905906 |
| **cg07955474** | -0.0076 | 0.262945331 | 0.269209212 | 0.278897053 | 0.409565727 | 0.135877454 |
| **cg07960624** | -0.01129 | 0.293567433 | 0.270853189 | 0.324777692 | 0.391313117 | 0.128393198 |
| **cg08548559** | -0.00837 | 0.190338448 | 0.185755602 | 0.359888815 | 0.384153789 | 0.239100657 |
| **cg08639339** | 0.00658 | 0.264639355 | 0.319104121 | 0.456895655 | 0.42115053 | 0.149754506 |
| **cg08857797** | 0.00833 | 0.171808414 | 0.191843106 | 0.243328041 | 0.231983911 | 0.101225363 |
| **cg09182678** | -0.00832 | 0.209034692 | 0.223952097 | 0.242288244 | 0.262941219 | 0.107668349 |
| **cg09349128** | -0.00741 | 0.103092152 | 0.079480677 | 0.164370599 | 0.158911174 | 0.070106395 |
| **cg09491962** | -0.0063 | 0.147812121 | 0.088293481 | 0.198507752 | 0.133548699 | 0.084405631 |
| **cg09607047** | 0.00638 | 0.062887347 | -0.000559202 | 0.132365372 | 0.085300173 | 0.041227961 |
| **cg09664445** | 0.00641 | 0.149093208 | 0.085493367 | 0.12805231 | 0.17781489 | 0.092911137 |
| **cg10092518** | -0.00932 | 0.044259271 | -0.014590858 | 0.111147597 | 0.102735193 | -0.029205933 |
| **cg10179300** | 0.01219 | 0.153317919 | 0.152794351 | 0.327101865 | 0.26153424 | 0.031599725 |
| **cg10192877** | 0.00535 | 0.115101867 | 0.05938464 | 0.050256231 | 0.082022148 | 0.035478701 |
| **cg10474597** | 0.00955 | 0.070656157 | 0.042478793 | 0.117069006 | 0.074538754 | 0.039388295 |
| **cg10508317** | -0.01221 | 0.019947122 | -0.039823801 | 0.160880466 | 0.150695907 | 0.074731587 |
| **cg10717869** | 0.00756 | 0.108360914 | 0.081218585 | 0.208126094 | 0.236500631 | 0.079366574 |
| **cg10919522** | -0.00804 | 0.172845048 | 0.196590006 | 0.267230375 | 0.162331888 | 0.095966037 |
| **cg11024682** | 0.00955 | 0.177229844 | 0.221060152 | 0.24238402 | 0.306253959 | 0.081619576 |
| **cg11152384** | -0.01011 | 0.289740529 | 0.249819547 | 0.377237493 | 0.414883202 | 0.116468789 |
| **cg11202345** | 0.01275 | 0.138489904 | 0.082975076 | 0.193292843 | 0.171758098 | 0.13696257 |
| **cg11261850** | 0.00967 | 0.190344068 | 0.218833768 | 0.316897505 | 0.333742321 | 0.072686004 |
| **cg11673687** | 0.00687 | 0.01616552 | -0.012865219 | 0.045837152 | 0.019448326 | 0.028457644 |
| **cg12001357** | -0.0082 | 0.23676697 | 0.209515813 | 0.451914574 | 0.416431571 | 0.127353306 |
| **cg12458003** | 0.00708 | 0.124248735 | 0.090090399 | 0.217587494 | 0.198782968 | 0.048922984 |
| **cg12484113** | 0.00637 | 0.093951223 | 0.08869643 | 0.122473233 | 0.108300543 | 0.077454083 |
| **cg12593793** | -0.0065 | 0.196171805 | 0.138566065 | 0.23644301 | 0.180507113 | 0.096438546 |
| **cg13028635** | 0.00538 | 0.128354525 | 0.121383834 | 0.194359323 | 0.20078105 | 0.07559707 |
| **cg13123009** | 0.00577 | 0.176271102 | 0.167484611 | 0.255101017 | 0.261163062 | 0.129934271 |
| **cg13134297** | -0.00867 | 0.137590989 | 0.093541737 | 0.116227313 | 0.111677286 | 0.096155259 |
| **cg13139542** | 0.00769 | 0.387299598 | 0.310663925 | 0.386648261 | 0.515878681 | 0.285251566 |
| **cg13274938** | 0.00621 | 0.136487472 | 0.095416407 | 0.133701045 | 0.109294305 | 0.061001554 |
| **cg13276570** | 0.01174 | 0.001861467 | -0.008847806 | 0.103195304 | 0.161708687 | 0.004413479 |
| **cg13305415** | 0.01006 | 0.170437396 | 0.165515177 | 0.168234958 | 0.172048199 | -0.003475021 |
| **cg13708645** | 0.00692 | 0.313722809 | 0.344731531 | 0.53973423 | 0.499880232 | 0.146195869 |
| **cg14017402** | 0.00776 | 0.456551708 | 0.461329431 | 0.578088112 | 0.366354387 | 0.195158056 |
| **cg14333542** | 0.00724 | 0.066882913 | 0.045627739 | 0.115282104 | 0.153040065 | 0.016012242 |
| **cg14352682** | 0.00594 | 0.140587458 | 0.051108547 | 0.09710293 | 0.049925668 | -0.000436105 |
| **cg14476101** | -0.01079 | 0.416081396 | 0.402075143 | 0.497033345 | 0.390539954 | 0.179797436 |
| **cg14509967** | 0.01193 | 0.01943632 | -0.058871434 | 0.223967173 | 0.191197277 | 0.04869942 |
| **cg14780837** | 0.00949 | 0.112988826 | 0.029019879 | 0.135583982 | 0.142366999 | 0.003586599 |
| **cg14870271** | 0.01068 | 0.159055746 | 0.15511543 | 0.217228096 | 0.250523081 | 0.139348755 |
| **cg15880704** | 0.0057 | 0.156231851 | 0.183007288 | 0.220275463 | 0.213029754 | 0.108535846 |
| **cg16611584** | 0.01325 | 0.402310114 | 0.38493603 | 0.545281548 | 0.593324083 | 0.153941246 |
| **cg16739178** | 0.00665 | 0.129401368 | 0.103539072 | 0.15770972 | 0.152191328 | 0.051373453 |
| **cg17058475** | -0.01265 | 0.036318349 | 0.04413234 | 0.111537341 | 0.092543993 | 0.003953912 |
| **cg17194270** | 0.0092 | 0.195583932 | 0.143125102 | 0.232975568 | 0.262352616 | 0.117362604 |
| **cg17501210** | -0.01267 | 0.038861637 | 0.023461103 | 0.240209684 | 0.368162875 | 0.02662751 |
| **cg17641710** | 0.00632 | 0.089073931 | 0.093217891 | 0.080553019 | 0.10380535 | 0.053692611 |
| **cg17738521** | -0.01255 | 0.177283936 | 0.099453812 | 0.074214456 | 0.137185169 | 0.089262748 |
| **cg17782974** | 0.01071 | 0.101750385 | 0.174182142 | 0.164299778 | 0.218044261 | 0.131718873 |
| **cg17836612** | 0.00881 | 0.202070089 | 0.179911157 | 0.340748855 | 0.300555169 | 0.085590717 |
| **cg17901584** | -0.01079 | 0.181112484 | 0.15579511 | 0.271442812 | 0.171551044 | 0.112608421 |
| **cg18091083** | 0.00996 | 0.096525666 | 0.028922417 | 0.428739902 | 0.53927999 | -0.021990109 |
| **cg18098839** | -0.01073 | 0.096873714 | 0.030498926 | 0.087587783 | 0.043726001 | -0.030770829 |
| **cg18181703** | -0.00743 | 0.115107682 | 0.041514978 | 0.287218889 | 0.265989693 | 0.01398787 |
| **cg18772573** | 0.00825 | 0.155831163 | 0.159949565 | 0.172771358 | 0.1573437 | 0.134954261 |
| **cg19017142** | -0.01037 | 0.040944554 | 0.044600574 | 0.128364563 | 0.156828728 | 0.03355355 |
| **cg19750657** | 0.01183 | 0.146330966 | 0.075195082 | 0.233656071 | 0.190836982 | 0.114051025 |
| **cg20496314** | 0.00899 | 0.219848836 | 0.20312654 | 0.284499155 | 0.17718494 | 0.06543972 |
| **cg20507228** | 0.00857 | 0.068002499 | 0.040129134 | 0.207054693 | 0.245971597 | 0.023289333 |
| **cg21429551** | -0.01343 | 0.375050493 | 0.358734425 | 0.506738057 | 0.469017457 | 0.112944566 |
| **cg21670987** | -0.011 | 0.290815091 | 0.268113692 | 0.4585512 | 0.577782974 | 0.097605367 |
| **cg21766592** | -0.01158 | 0.145440377 | 0.15950758 | 0.246377426 | 0.190018274 | 0.066189391 |
| **cg22012981** | 0.00688 | 0.071776003 | 0.079971644 | 0.031087641 | 0.126920207 | 0.042408928 |
| **cg22650271** | 0.00764 | 0.040726238 | -0.022735505 | 0.069552418 | 0.123445995 | 0.01827102 |
| **cg22713958** | 0.00677 | 0.182962711 | 0.149135065 | 0.297978514 | 0.227462665 | 0.105877374 |
| **cg22875823** | 0.00923 | 0.218154241 | 0.241150442 | 0.329137257 | 0.292444259 | 0.122064772 |
| **cg22950899** | 0.0087 | 0.150994947 | 0.129555037 | 0.236518796 | 0.303398836 | 0.058142096 |
| **cg23172671** | 0.00857 | 0.011485411 | 0.104100862 | 0.113991213 | 0.18553046 | -0.002372929 |
| **cg23813257** | -0.00551 | 0.304867078 | 0.306911907 | 0.351492701 | 0.327044418 | 0.136028722 |
| **cg23998749** | 0.00704 | 0.136615331 | 0.072846934 | 0.091125943 | 0.184501052 | 0.052236701 |
| **cg24145109** | 0.01057 | 0.445401948 | 0.4177352 | 0.585171322 | 0.507263003 | 0.229488225 |
| **cg24531955** | -0.00863 | 0.264668996 | 0.22513148 | 0.296307258 | 0.284810026 | 0.186668642 |
| **cg24678869** | 0.00557 | 0.079609465 | 0.083421676 | 0.129234762 | 0.18822151 | 0.043618632 |
| **cg25178683** | 0.00946 | 0.213337628 | 0.215975427 | 0.358844842 | 0.34914817 | 0.144030352 |
| **cg25217710** | 0.00551 | 0.140094168 | 0.160942029 | 0.182012595 | 0.20648137 | 0.070274436 |
| **cg25392060** | 0.00598 | 0.253438132 | 0.201821272 | 0.22823457 | 0.251420287 | 0.141936811 |
| **cg25649826** | 0.00699 | 0.21517196 | 0.151074329 | 0.180177303 | 0.19846994 | 0.138730175 |
| **cg26191447** | -0.01107 | 0.270554289 | 0.252946374 | 0.335824407 | 0.530161419 | 0.09465428 |
| **cg26361535** | 0.00952 | 0.290594756 | 0.154539289 | 0.297775886 | 0.366391437 | 0.037793556 |
| **cg26403843** | 0.0134 | 0.434726321 | 0.385453017 | 0.475509285 | 0.468813114 | 0.22230044 |
| **cg26470501** | -0.00758 | 0.265139014 | 0.278083678 | 0.327435596 | 0.315728422 | 0.178495385 |
| **cg26651978** | -0.0071 | 0.09372085 | 0.052713153 | 0.189322604 | 0.23700832 | 0.021189951 |
| **cg26800893** | -0.00763 | 0.098282021 | 0.062456779 | 0.089925103 | 0.031347787 | 0.008586107 |
| **cg26894079** | -0.00886 | 0.127245807 | 0.162829675 | 0.272082367 | 0.284660462 | 0.052144571 |
| **cg26950531** | -0.01174 | 0.137886405 | 0.068836444 | 0.147984611 | 0.166115479 | 0.029271656 |
| **cg26955383** | 0.00722 | 0.096600036 | 0.108589752 | 0.138309586 | 0.221971672 | 0.105137889 |
| **cg27115863** | -0.01042 | 0.237225081 | 0.193199486 | 0.344065552 | 0.323873968 | 0.066191166 |
| **cg27243685** | 0.01107 | 0.059845126 | 0.073311653 | 0.10126109 | 0.142719193 | 0.056581208 |
| **cg27394566** | 0.00926 | 0.166175035 | 0.185857366 | 0.228499773 | 0.228225507 | 0.103122486 |
| **cg27470213** | -0.01108 | 0.125351028 | 0.06783173 | 0.255231673 | 0.184937416 | 0.092576534 |
| **cg27637521** | -0.01198 | -0.031584297 | -0.056468914 | 0.058010094 | 0.086030499 | -0.017758064 |

**Supplementary table 2**. Table of Mendelian Randomisation type analyses to assess whether there is a causal association from BMI to each of th2 135 CpG sites used to construct the methylation score.

| **CpG** | **Estimate (Lower CI, Upper CI, P-value)** | | | | |
| --- | --- | --- | --- | --- | --- |
|  | **Cord** | **Childhood** | **Adolescence** | **Middle-age** | **Antenatal** |
| **cg01455178** | -0.0175 (-0.0849, 0.0501, p=6.12E-01) | 0.0119 (-0.0531, 0.0768, p=7.20E-01) | 0.0249 (-0.0402, 0.0899, p=4.53E-01) | -0.0319 (-0.0983, 0.0347, p=3.48E-01) | -0.0287 (-0.0956, 0.0384, p=4.01E-01) |
| **cg03725309** | -0.002 (-0.0695, 0.0655, p=9.54E-01) | -0.0489 (-0.1135, 0.0161, p=1.40E-01) | 0.0086 (-0.0565, 0.0737, p=7.95E-01) | -0.0029 (-0.0695, 0.0637, p=9.31E-01) | 0.0371 (-0.03, 0.1039, p=2.79E-01) |
| **cg08639339** | -0.0054 (-0.0729, 0.0621, p=8.75E-01) | -0.0354 (-0.1001, 0.0296, p=2.86E-01) | -0.0114 (-0.0765, 0.0537, p=7.31E-01) | -0.0372 (-0.1035, 0.0295, p=2.74E-01) | -0.0195 (-0.0865, 0.0476, p=5.68E-01) |
| **cg10092518** | -0.0161 (-0.0835, 0.0514, p=6.40E-01) | -0.0151 (-0.08, 0.0499, p=6.48E-01) | -0.0189 (-0.0839, 0.0463, p=5.70E-01) | -0.0367 (-0.103, 0.03, p=2.80E-01) | -0.0049 (-0.0719, 0.0622, p=8.86E-01) |
| **cg10717869** | 0.0038 (-0.0637, 0.0713, p=9.12E-01) | -0.0311 (-0.0959, 0.0339, p=3.48E-01) | 0.0066 (-0.0585, 0.0717, p=8.42E-01) | 0.028 (-0.0386, 0.0944, p=4.10E-01) | -0.0416 (-0.1083, 0.0255, p=2.24E-01) |
| **cg11673687** | 0.0318 (-0.0358, 0.099, p=3.57E-01) | 0.0332 (-0.0319, 0.0979, p=3.17E-01) | 0.0208 (-0.0443, 0.0858, p=5.31E-01) | -0.0561 (-0.1223, 0.0105, p=9.85E-02) | 0.0049 (-0.0621, 0.072, p=8.85E-01) |
| **cg12458003** | 0.0671 (-0.0004, 0.134, p=5.14E-02) | -0.0025 (-0.0674, 0.0625, p=9.41E-01) | 0.0138 (-0.0513, 0.0788, p=6.78E-01) | 0.0024 (-0.0642, 0.069, p=9.43E-01) | 0.0355 (-0.0317, 0.1023, p=3.00E-01) |
| **cg12484113** | -0.0265 (-0.0938, 0.0411, p=4.43E-01) | -0.0406 (-0.1053, 0.0244, p=2.20E-01) | -0.0317 (-0.0966, 0.0335, p=3.40E-01) | 0.041 (-0.0256, 0.1073, p=2.28E-01) | 0.032 (-0.0351, 0.0989, p=3.50E-01) |
| **cg12593793** | -0.0414 (-0.1085, 0.0262, p=2.30E-01) | -0.0501 (-0.1147, 0.0149, p=1.30E-01) | -0.034 (-0.0989, 0.0311, p=3.06E-01) | -0.0099 (-0.0764, 0.0567, p=7.71E-01) | 0.0226 (-0.0445, 0.0895, p=5.09E-01) |
| **cg14476101** | 0.0156 (-0.0519, 0.083, p=6.50E-01) | -0.008 (-0.0729, 0.057, p=8.09E-01) | -0.0126 (-0.0777, 0.0525, p=7.04E-01) | -0.0169 (-0.0834, 0.0498, p=6.20E-01) | -0.0089 (-0.0759, 0.0582, p=7.95E-01) |
| **cg17901584** | 0.0059 (-0.0616, 0.0734, p=8.64E-01) | 0.0166 (-0.0484, 0.0815, p=6.16E-01) | -0.013 (-0.0781, 0.0521, p=6.95E-01) | 0.0029 (-0.0637, 0.0695, p=9.31E-01) | 0.0032 (-0.0638, 0.0703, p=9.25E-01) |
| **cg23172671** | -0.0219 (-0.0893, 0.0456, p=5.24E-01) | -0.0351 (-0.0998, 0.0299, p=2.90E-01) | 0.0147 (-0.0505, 0.0797, p=6.59E-01) | 0.04 (-0.0267, 0.1063, p=2.40E-01) | 0.0164 (-0.0507, 0.0834, p=6.31E-01) |
| **cg23998749** | -0.0161 (-0.0835, 0.0514, p=6.40E-01) | -0.0068 (-0.0718, 0.0581, p=8.37E-01) | -0.0145 (-0.0795, 0.0506, p=6.62E-01) | 0.0485 (-0.0182, 0.1147, p=1.54E-01) | -0.0459 (-0.1126, 0.0212, p=1.80E-01) |
| **cg24145109** | -0.0004 (-0.0679, 0.0671, p=9.91E-01) | -0.0165 (-0.0814, 0.0485, p=6.19E-01) | -0.0046 (-0.0697, 0.0605, p=8.90E-01) | 0.0075 (-0.0591, 0.074, p=8.26E-01) | 0.0089 (-0.0582, 0.0759, p=7.95E-01) |
| **cg24678869** | -0.0105 (-0.0779, 0.057, p=7.60E-01) | -0.019 (-0.0838, 0.046, p=5.67E-01) | -0.0066 (-0.0716, 0.0586, p=8.44E-01) | -0.0221 (-0.0885, 0.0446, p=5.16E-01) | 0.0034 (-0.0637, 0.0704, p=9.22E-01) |
| **cg25217710** | -0.029 (-0.0963, 0.0386, p=4.00E-01) | -0.0443 (-0.109, 0.0207, p=1.81E-01) | -0.0025 (-0.0676, 0.0626, p=9.39E-01) | 0.0063 (-0.0603, 0.0729, p=8.53E-01) | 0.0074 (-0.0597, 0.0744, p=8.29E-01) |
| **cg04011474** | 0.0613 (-0.0062, 0.1283, p=7.49E-02) | -0.0458 (-0.1104, 0.0192, p=1.67E-01) | -0.0376 (-0.1025, 0.0275, p=2.57E-01) | -0.0155 (-0.082, 0.0511, p=6.49E-01) | 0.0005 (-0.0665, 0.0676, p=9.88E-01) |
| **cg04286697** | 0.048 (-0.0195, 0.1151, p=1.64E-01) | -0.0396 (-0.1043, 0.0255, p=2.33E-01) | 0.0124 (-0.0528, 0.0774, p=7.10E-01) | -0.0126 (-0.0791, 0.0541, p=7.12E-01) | -0.0018 (-0.0688, 0.0653, p=9.58E-01) |
| **cg06876354** | 0.0443 (-0.0232, 0.1115, p=1.98E-01) | -0.0021 (-0.067, 0.0629, p=9.50E-01) | -0.0036 (-0.0687, 0.0615, p=9.13E-01) | 0.053 (-0.0136, 0.1191, p=1.19E-01) | 0.0717 (0.0047, 0.1381, p=3.60E-02) |
| **cg12001357** | -0.0151 (-0.0825, 0.0525, p=6.62E-01) | 0.0137 (-0.0513, 0.0786, p=6.80E-01) | -0.0306 (-0.0955, 0.0346, p=3.57E-01) | -0.0017 (-0.0683, 0.0649, p=9.59E-01) | -0.0124 (-0.0794, 0.0547, p=7.18E-01) |
| **cg13139542** | -0.038 (-0.1052, 0.0296, p=2.71E-01) | -0.0287 (-0.0935, 0.0363, p=3.87E-01) | -0.0106 (-0.0756, 0.0546, p=7.50E-01) | -0.0058 (-0.0723, 0.0608, p=8.65E-01) | -0.0083 (-0.0753, 0.0588, p=8.09E-01) |
| **cg14017402** | -0.0244 (-0.0917, 0.0432, p=4.79E-01) | -0.0461 (-0.1107, 0.0189, p=1.64E-01) | -0.0047 (-0.0698, 0.0604, p=8.88E-01) | 0.0514 (-0.0152, 0.1176, p=1.31E-01) | 0.0154 (-0.0517, 0.0824, p=6.53E-01) |
| **cg19017142** | 0.0044 (-0.0631, 0.0719, p=8.97E-01) | 0.003 (-0.0619, 0.068, p=9.27E-01) | -0.0338 (-0.0986, 0.0314, p=3.10E-01) | -0.042 (-0.1083, 0.0246, p=2.16E-01) | 0.0357 (-0.0314, 0.1025, p=2.97E-01) |
| **cg26191447** | -0.0218 (-0.0891, 0.0458, p=5.27E-01) | -0.031 (-0.0958, 0.034, p=3.50E-01) | 0.0277 (-0.0375, 0.0926, p=4.05E-01) | 0.0451 (-0.0215, 0.1114, p=1.84E-01) | 0.007 (-0.0601, 0.074, p=8.39E-01) |
| **cg00108715** | 0.0622 (-0.0053, 0.1291, p=7.10E-02) | -0.0262 (-0.091, 0.0388, p=4.30E-01) | 0.0015 (-0.0636, 0.0666, p=9.64E-01) | -0.0058 (-0.0723, 0.0608, p=8.65E-01) | 0.0606 (-0.0065, 0.1271, p=7.65E-02) |
| **cg01368219** | 0.0042 (-0.0633, 0.0717, p=9.03E-01) | -0.0122 (-0.0771, 0.0528, p=7.14E-01) | 0.0217 (-0.0434, 0.0867, p=5.13E-01) | 0.0332 (-0.0335, 0.0995, p=3.29E-01) | -0.0404 (-0.1071, 0.0267, p=2.38E-01) |
| **cg01526748** | -0.0372 (-0.1044, 0.0304, p=2.81E-01) | 0.0212 (-0.0438, 0.086, p=5.23E-01) | 0.0416 (-0.0235, 0.1064, p=2.11E-01) | 0.0623 (-0.0043, 0.1284, p=6.66E-02) | 0.0414 (-0.0258, 0.1081, p=2.27E-01) |
| **cg01671681** | 0.0007 (-0.0668, 0.0682, p=9.83E-01) | -0.0278 (-0.0926, 0.0372, p=4.01E-01) | -0.0639 (-0.1285, 0.0012, p=5.42E-02) | -0.0333 (-0.0997, 0.0333, p=3.27E-01) | -0.0186 (-0.0856, 0.0485, p=5.87E-01) |
| **cg07730360** | 0.0225 (-0.045, 0.0899, p=5.13E-01) | -0.0325 (-0.0972, 0.0325, p=3.27E-01) | 0.0642 (-0.0009, 0.1288, p=5.31E-02) | 0.0141 (-0.0525, 0.0806, p=6.78E-01) | 0.0519 (-0.0152, 0.1186, p=1.29E-01) |
| **cg17641710** | -0.0145 (-0.0819, 0.0531, p=6.74E-01) | -0.0088 (-0.0737, 0.0562, p=7.91E-01) | -0.0451 (-0.1099, 0.0201, p=1.75E-01) | 0.0185 (-0.0481, 0.085, p=5.87E-01) | -0.0099 (-0.0769, 0.0572, p=7.72E-01) |
| **cg18098839** | 0.0325 (-0.0351, 0.0998, p=3.46E-01) | -0.0122 (-0.0771, 0.0528, p=7.13E-01) | -0.0394 (-0.1042, 0.0258, p=2.36E-01) | 0.0199 (-0.0468, 0.0863, p=5.59E-01) | 0.0286 (-0.0386, 0.0954, p=4.04E-01) |
| **cg22012981** | 0.033 (-0.0346, 0.1002, p=3.39E-01) | -0.0025 (-0.0674, 0.0625, p=9.41E-01) | 0.031 (-0.0342, 0.0959, p=3.52E-01) | -0.0037 (-0.0702, 0.0629, p=9.14E-01) | -0.0538 (-0.1204, 0.0133, p=1.16E-01) |
| **cg05119988** | -0.0109 (-0.0783, 0.0566, p=7.52E-01) | 0.0146 (-0.0504, 0.0794, p=6.61E-01) | -0.0603 (-0.1249, 0.0048, p=6.95E-02) | 0.052 (-0.0146, 0.1182, p=1.26E-01) | 0.0033 (-0.0638, 0.0703, p=9.23E-01) |
| **cg06690548** | 0.0267 (-0.0409, 0.094, p=4.39E-01) | 0.0357 (-0.0294, 0.1004, p=2.82E-01) | 0.0129 (-0.0522, 0.0779, p=6.98E-01) | 0.0047 (-0.0619, 0.0712, p=8.91E-01) | -0.0278 (-0.0947, 0.0393, p=4.17E-01) |
| **cg07094298** | 0.0037 (-0.0638, 0.0712, p=9.14E-01) | 0.0107 (-0.0543, 0.0756, p=7.48E-01) | -0.0233 (-0.0882, 0.0419, p=4.84E-01) | 0.0027 (-0.0639, 0.0693, p=9.36E-01) | -0.0374 (-0.1042, 0.0297, p=2.74E-01) |
| **cg02286155** | 0.0358 (-0.0318, 0.103, p=2.99E-01) | 0.0222 (-0.0428, 0.087, p=5.03E-01) | 0.0802 (0.0152, 0.1446, p=1.57E-02) | -0.0213 (-0.0878, 0.0453, p=5.31E-01) | -0.0213 (-0.0882, 0.0459, p=5.35E-01) |
| **cg04483863** | -0.0203 (-0.0877, 0.0473, p=5.56E-01) | 0.0092 (-0.0558, 0.0741, p=7.82E-01) | 0.0709 (0.0059, 0.1354, p=3.27E-02) | -0.0414 (-0.1077, 0.0252, p=2.23E-01) | -0.0314 (-0.0982, 0.0357, p=3.59E-01) |
| **cg10179300** | -0.0737 (-0.1405, -0.0063, p=3.22E-02) | -0.0264 (-0.0912, 0.0386, p=4.26E-01) | 0.0223 (-0.0428, 0.0873, p=5.02E-01) | 0.0176 (-0.049, 0.0841, p=6.04E-01) | -0.0142 (-0.0812, 0.0529, p=6.79E-01) |
| **cg13276570** | 0.0131 (-0.0545, 0.0805, p=7.05E-01) | -0.0378 (-0.1025, 0.0272, p=2.55E-01) | 0.0029 (-0.0622, 0.068, p=9.30E-01) | 0.0188 (-0.0478, 0.0853, p=5.80E-01) | -0.0671 (-0.1335, 0, p=4.99E-02) |
| **cg13305415** | 0.0025 (-0.065, 0.07, p=9.41E-01) | -0.0034 (-0.0684, 0.0615, p=9.18E-01) | -0.0227 (-0.0877, 0.0425, p=4.95E-01) | -0.0132 (-0.0797, 0.0534, p=6.98E-01) | 0.0117 (-0.0554, 0.0787, p=7.33E-01) |
| **cg26403843** | -0.0602 (-0.1272, 0.0073, p=8.05E-02) | -0.047 (-0.1117, 0.018, p=1.56E-01) | -0.0317 (-0.0966, 0.0334, p=3.40E-01) | -0.0013 (-0.0678, 0.0653, p=9.70E-01) | -0.0405 (-0.1072, 0.0266, p=2.37E-01) |
| **cg13123009** | -0.0517 (-0.1188, 0.0158, p=1.33E-01) | -0.0267 (-0.0915, 0.0383, p=4.21E-01) | -0.0093 (-0.0744, 0.0558, p=7.79E-01) | 0.0283 (-0.0383, 0.0947, p=4.05E-01) | -0.0122 (-0.0792, 0.0549, p=7.21E-01) |
| **cg14352682** | 0.0269 (-0.0406, 0.0942, p=4.35E-01) | 0.0043 (-0.0607, 0.0692, p=8.98E-01) | -0.0155 (-0.0805, 0.0496, p=6.41E-01) | 0.0114 (-0.0552, 0.0779, p=7.37E-01) | 0.0316 (-0.0355, 0.0984, p=3.56E-01) |
| **cg17501210** | 0.0182 (-0.0493, 0.0856, p=5.97E-01) | 0.0163 (-0.0487, 0.0811, p=6.24E-01) | -0.0166 (-0.0816, 0.0486, p=6.18E-01) | -0.047 (-0.1132, 0.0196, p=1.67E-01) | -0.0143 (-0.0813, 0.0528, p=6.76E-01) |
| **cg17738521** | -0.0002 (-0.0677, 0.0673, p=9.95E-01) | -0.0051 (-0.07, 0.0599, p=8.77E-01) | 0.0311 (-0.0341, 0.096, p=3.50E-01) | 0.0233 (-0.0433, 0.0898, p=4.93E-01) | 0.0164 (-0.0507, 0.0834, p=6.31E-01) |
| **cg22875823** | 0.0128 (-0.0547, 0.0802, p=7.11E-01) | -0.0105 (-0.0754, 0.0545, p=7.52E-01) | 0.0321 (-0.033, 0.097, p=3.33E-01) | -0.0051 (-0.0717, 0.0615, p=8.80E-01) | 0.0432 (-0.0239, 0.1099, p=2.07E-01) |
| **cg04816311** | 0.0497 (-0.0178, 0.1168, p=1.49E-01) | 0.0088 (-0.0561, 0.0737, p=7.90E-01) | 0.0302 (-0.035, 0.0951, p=3.64E-01) | -0.056 (-0.1221, 0.0106, p=9.93E-02) | 0.0038 (-0.0633, 0.0708, p=9.12E-01) |
| **cg13134297** | -0.0374 (-0.1046, 0.0302, p=2.78E-01) | 0.0185 (-0.0465, 0.0834, p=5.77E-01) | -0.0124 (-0.0774, 0.0527, p=7.09E-01) | -0.0221 (-0.0885, 0.0445, p=5.16E-01) | -0.0138 (-0.0808, 0.0533, p=6.86E-01) |
| **cg21429551** | -0.0246 (-0.092, 0.0429, p=4.75E-01) | 0.0053 (-0.0597, 0.0702, p=8.74E-01) | -0.0316 (-0.0965, 0.0336, p=3.42E-01) | 0.0087 (-0.058, 0.0752, p=7.99E-01) | -0.0267 (-0.0935, 0.0405, p=4.36E-01) |
| **cg02571142** | -0.0026 (-0.07, 0.0649, p=9.41E-01) | -0.0789 (-0.1431, -0.014, p=1.73E-02) | -0.0232 (-0.0881, 0.042, p=4.86E-01) | -0.0322 (-0.0986, 0.0344, p=3.43E-01) | 0.0031 (-0.064, 0.0701, p=9.28E-01) |
| **cg07960624** | 0.0487 (-0.0188, 0.1158, p=1.57E-01) | -0.0088 (-0.0737, 0.0562, p=7.91E-01) | -0.0241 (-0.089, 0.0411, p=4.69E-01) | -0.0149 (-0.0814, 0.0517, p=6.61E-01) | -0.0064 (-0.0734, 0.0607, p=8.52E-01) |
| **cg24531955** | 0.0179 (-0.0497, 0.0852, p=6.04E-01) | -0.0089 (-0.0738, 0.0561, p=7.89E-01) | -0.0288 (-0.0937, 0.0364, p=3.87E-01) | 0.0113 (-0.0554, 0.0778, p=7.41E-01) | 0.0039 (-0.0632, 0.0709, p=9.10E-01) |
| **cg25392060** | -0.0084 (-0.0758, 0.0591, p=8.08E-01) | -0.0467 (-0.1113, 0.0183, p=1.59E-01) | -0.0282 (-0.0931, 0.037, p=3.96E-01) | -0.0149 (-0.0814, 0.0517, p=6.61E-01) | -0.0033 (-0.0703, 0.0637, p=9.23E-01) |
| **cg26361535** | 0.0128 (-0.0547, 0.0802, p=7.11E-01) | -0.0395 (-0.1041, 0.0256, p=2.34E-01) | 0.0598 (-0.0054, 0.1244, p=7.21E-02) | -0.0239 (-0.0904, 0.0427, p=4.82E-01) | -0.0698 (-0.1362, -0.0027, p=4.14E-02) |
| **cg02716826** | 0.0199 (-0.0476, 0.0873, p=5.63E-01) | -0.0274 (-0.0922, 0.0376, p=4.08E-01) | 0.0038 (-0.0613, 0.0688, p=9.10E-01) | 0.04 (-0.0266, 0.1063, p=2.39E-01) | 0.0036 (-0.0635, 0.0706, p=9.17E-01) |
| **cg07504977** | -0.0233 (-0.0907, 0.0442, p=4.99E-01) | -0.0779 (-0.1421, -0.013, p=1.88E-02) | -0.0357 (-0.1006, 0.0294, p=2.82E-01) | 0.0048 (-0.0618, 0.0714, p=8.87E-01) | -0.0141 (-0.0811, 0.053, p=6.80E-01) |
| **cg09491962** | -0.0015 (-0.069, 0.0659, p=9.64E-01) | -0.0218 (-0.0867, 0.0432, p=5.10E-01) | -0.0155 (-0.0805, 0.0497, p=6.42E-01) | -0.0204 (-0.0869, 0.0462, p=5.49E-01) | 0.0177 (-0.0494, 0.0846, p=6.06E-01) |
| **cg14333542** | 0.0114 (-0.0561, 0.0789, p=7.40E-01) | -0.0222 (-0.087, 0.0428, p=5.04E-01) | -0.0098 (-0.0749, 0.0553, p=7.67E-01) | -0.0678 (-0.1337, -0.0012, p=4.61E-02) | 0.0043 (-0.0627, 0.0713, p=9.00E-01) |
| **cg15880704** | -0.0074 (-0.0748, 0.0601, p=8.30E-01) | 0.0397 (-0.0253, 0.1044, p=2.31E-01) | 0.0179 (-0.0472, 0.0829, p=5.90E-01) | -0.0243 (-0.0907, 0.0424, p=4.75E-01) | -0.0465 (-0.1132, 0.0206, p=1.74E-01) |
| **cg17782974** | 0.0017 (-0.0658, 0.0691, p=9.62E-01) | 0.0054 (-0.0596, 0.0703, p=8.71E-01) | -0.0339 (-0.0988, 0.0313, p=3.08E-01) | 0.0118 (-0.0548, 0.0783, p=7.29E-01) | -0.0399 (-0.1066, 0.0272, p=2.44E-01) |
| **cg26955383** | 0.0149 (-0.0526, 0.0823, p=6.66E-01) | -0.0036 (-0.0685, 0.0614, p=9.14E-01) | -0.0114 (-0.0764, 0.0538, p=7.32E-01) | 0.0393 (-0.0273, 0.1056, p=2.47E-01) | 0.0109 (-0.0562, 0.0779, p=7.51E-01) |
| **cg00574958** | -0.0229 (-0.0903, 0.0446, p=5.06E-01) | 0.0012 (-0.0638, 0.0661, p=9.72E-01) | 0.0064 (-0.0587, 0.0715, p=8.47E-01) | 0.0239 (-0.0428, 0.0903, p=4.83E-01) | 0.002 (-0.0651, 0.069, p=9.53E-01) |
| **cg11152384** | 0.0543 (-0.0132, 0.1214, p=1.15E-01) | -0.0415 (-0.1062, 0.0235, p=2.10E-01) | 0.0055 (-0.0596, 0.0706, p=8.69E-01) | -0.0422 (-0.1085, 0.0244, p=2.14E-01) | -0.0152 (-0.0822, 0.0519, p=6.57E-01) |
| **cg11261850** | -0.0223 (-0.0896, 0.0453, p=5.18E-01) | 0.0153 (-0.0497, 0.0802, p=6.44E-01) | -0.0042 (-0.0693, 0.0609, p=8.99E-01) | -0.0275 (-0.0939, 0.0391, p=4.18E-01) | -0.0246 (-0.0915, 0.0425, p=4.72E-01) |
| **cg13028635** | 0.0409 (-0.0267, 0.1081, p=2.36E-01) | 0.0323 (-0.0327, 0.0971, p=3.30E-01) | 0.0535 (-0.0116, 0.1182, p=1.07E-01) | -0.0051 (-0.0716, 0.0615, p=8.81E-01) | -0.0197 (-0.0866, 0.0474, p=5.65E-01) |
| **cg17058475** | 0.0207 (-0.0468, 0.0881, p=5.48E-01) | -0.0221 (-0.0869, 0.0429, p=5.05E-01) | -0.0135 (-0.0786, 0.0516, p=6.84E-01) | 0.0073 (-0.0593, 0.0738, p=8.30E-01) | 0.0092 (-0.0579, 0.0762, p=7.89E-01) |
| **cg26800893** | 0.0031 (-0.0644, 0.0706, p=9.28E-01) | -0.0563 (-0.1208, 0.0087, p=8.95E-02) | 0.0226 (-0.0425, 0.0876, p=4.96E-01) | 0.0073 (-0.0594, 0.0738, p=8.31E-01) | 0.0265 (-0.0406, 0.0934, p=4.38E-01) |
| **cg26894079** | 0.0034 (-0.0641, 0.0709, p=9.21E-01) | -0.0074 (-0.0724, 0.0575, p=8.23E-01) | 0.017 (-0.0482, 0.082, p=6.10E-01) | 0.0111 (-0.0556, 0.0776, p=7.45E-01) | 0.0104 (-0.0567, 0.0774, p=7.62E-01) |
| **cg13708645** | -0.0581 (-0.1251, 0.0094, p=9.16E-02) | 0.0159 (-0.0491, 0.0808, p=6.31E-01) | 0.0162 (-0.0489, 0.0812, p=6.25E-01) | -0.051 (-0.1172, 0.0156, p=1.33E-01) | -0.0258 (-0.0927, 0.0413, p=4.51E-01) |
| **cg10474597** | -0.0266 (-0.0939, 0.041, p=4.40E-01) | -0.008 (-0.0729, 0.057, p=8.10E-01) | 0.0316 (-0.0335, 0.0965, p=3.41E-01) | 0.004 (-0.0626, 0.0706, p=9.05E-01) | 0.0153 (-0.0518, 0.0823, p=6.55E-01) |
| **cg19750657** | 0.0137 (-0.0538, 0.0811, p=6.91E-01) | 0.0035 (-0.0615, 0.0684, p=9.16E-01) | -0.0272 (-0.0921, 0.038, p=4.14E-01) | 0.0085 (-0.0581, 0.0751, p=8.02E-01) | 0.0437 (-0.0234, 0.1104, p=2.02E-01) |
| **cg10919522** | 0.0031 (-0.0644, 0.0705, p=9.29E-01) | 0.006 (-0.059, 0.0709, p=8.57E-01) | -0.0514 (-0.1161, 0.0138, p=1.22E-01) | 0.0275 (-0.0392, 0.0939, p=4.19E-01) | -0.0267 (-0.0936, 0.0404, p=4.36E-01) |
| **cg27394566** | -0.0094 (-0.0769, 0.0581, p=7.84E-01) | -0.0094 (-0.0743, 0.0556, p=7.77E-01) | 0.049 (-0.0161, 0.1138, p=1.40E-01) | -0.0574 (-0.1235, 0.0092, p=9.13E-02) | -0.0419 (-0.1087, 0.0252, p=2.21E-01) |
| **cg06192883** | -0.0199 (-0.0872, 0.0477, p=5.64E-01) | -0.0181 (-0.0829, 0.047, p=5.86E-01) | -0.0045 (-0.0696, 0.0606, p=8.92E-01) | 0.0566 (-0.01, 0.1228, p=9.56E-02) | 0.0071 (-0.0599, 0.0742, p=8.35E-01) |
| **cg07037944** | 0.0099 (-0.0576, 0.0773, p=7.74E-01) | -0.0519 (-0.1165, 0.0131, p=1.17E-01) | -0.0227 (-0.0877, 0.0424, p=4.94E-01) | 0.0163 (-0.0503, 0.0828, p=6.31E-01) | 0.0302 (-0.0369, 0.0971, p=3.77E-01) |
| **cg07814318** | -0.051 (-0.1181, 0.0166, p=1.39E-01) | -0.0032 (-0.0682, 0.0617, p=9.23E-01) | 0.0216 (-0.0436, 0.0865, p=5.16E-01) | -0.0115 (-0.078, 0.0551, p=7.35E-01) | -0.003 (-0.07, 0.064, p=9.30E-01) |
| **cg20507228** | -0.0122 (-0.0796, 0.0553, p=7.23E-01) | 0.0106 (-0.0544, 0.0755, p=7.50E-01) | -0.0162 (-0.0813, 0.0489, p=6.25E-01) | 0.013 (-0.0536, 0.0795, p=7.01E-01) | -0.0043 (-0.0713, 0.0628, p=9.00E-01) |
| **cg21670987** | 0.0336 (-0.0339, 0.1009, p=3.29E-01) | 0.0512 (-0.0138, 0.1158, p=1.23E-01) | 0.0127 (-0.0524, 0.0778, p=7.02E-01) | -0.0162 (-0.0827, 0.0505, p=6.34E-01) | 0.002 (-0.0651, 0.069, p=9.53E-01) |
| **cg01243823** | 0.0715 (0.0041, 0.1384, p=3.77E-02) | -0.0431 (-0.1077, 0.0219, p=1.94E-01) | -0.0592 (-0.1238, 0.006, p=7.49E-02) | -0.0151 (-0.0816, 0.0516, p=6.58E-01) | -0.0231 (-0.09, 0.044, p=5.00E-01) |
| **cg03500056** | -0.0296 (-0.0968, 0.038, p=3.91E-01) | 0.0171 (-0.0479, 0.0819, p=6.07E-01) | -0.0062 (-0.0713, 0.0589, p=8.52E-01) | -0.0047 (-0.0712, 0.0619, p=8.91E-01) | 0.0223 (-0.0449, 0.0892, p=5.16E-01) |
| **cg06946797** | 0.0266 (-0.0409, 0.094, p=4.39E-01) | -0.0166 (-0.0814, 0.0484, p=6.17E-01) | -0.042 (-0.1068, 0.0232, p=2.07E-01) | -0.0496 (-0.1158, 0.017, p=1.44E-01) | 0.024 (-0.0431, 0.0909, p=4.83E-01) |
| **cg07021906** | -0.0225 (-0.0899, 0.045, p=5.14E-01) | -0.0273 (-0.0921, 0.0377, p=4.10E-01) | 0.035 (-0.0302, 0.0998, p=2.93E-01) | 0.0079 (-0.0588, 0.0744, p=8.17E-01) | -0.0069 (-0.0739, 0.0602, p=8.41E-01) |
| **cg07955474** | -0.0392 (-0.1064, 0.0283, p=2.55E-01) | -0.0505 (-0.115, 0.0145, p=1.28E-01) | -0.0323 (-0.0972, 0.0329, p=3.31E-01) | -0.0086 (-0.0751, 0.058, p=8.00E-01) | -0.004 (-0.0711, 0.063, p=9.06E-01) |
| **cg09607047** | -0.04 (-0.1072, 0.0276, p=2.46E-01) | 0.0164 (-0.0486, 0.0813, p=6.21E-01) | 0.0191 (-0.046, 0.0841, p=5.65E-01) | 0.0218 (-0.0448, 0.0883, p=5.21E-01) | 0.0055 (-0.0615, 0.0725, p=8.72E-01) |
| **cg16739178** | 0.0086 (-0.059, 0.076, p=8.04E-01) | -0.0124 (-0.0772, 0.0526, p=7.10E-01) | 0.0451 (-0.02, 0.1099, p=1.75E-01) | -0.0388 (-0.1051, 0.0278, p=2.54E-01) | -0.0007 (-0.0677, 0.0664, p=9.84E-01) |
| **cg23813257** | 0.0302 (-0.0373, 0.0975, p=3.80E-01) | 0.0155 (-0.0495, 0.0804, p=6.40E-01) | 0.013 (-0.0522, 0.078, p=6.96E-01) | 0.0248 (-0.0419, 0.0912, p=4.66E-01) | -0.0176 (-0.0846, 0.0495, p=6.07E-01) |
| **cg01130991** | -0.0358 (-0.103, 0.0318, p=2.99E-01) | -0.0047 (-0.0697, 0.0602, p=8.86E-01) | 0.0221 (-0.043, 0.0871, p=5.06E-01) | -0.0206 (-0.0871, 0.046, p=5.44E-01) | -0.0037 (-0.0707, 0.0633, p=9.14E-01) |
| **cg01597398** | -0.005 (-0.0724, 0.0625, p=8.86E-01) | -0.0063 (-0.0713, 0.0586, p=8.49E-01) | -0.0435 (-0.1083, 0.0216, p=1.90E-01) | -0.0145 (-0.081, 0.0521, p=6.69E-01) | 0.0075 (-0.0596, 0.0745, p=8.26E-01) |
| **cg01798813** | -0.0286 (-0.0959, 0.0389, p=4.06E-01) | -0.0057 (-0.0707, 0.0592, p=8.63E-01) | 0.0094 (-0.0557, 0.0744, p=7.77E-01) | -0.009 (-0.0756, 0.0576, p=7.90E-01) | -0.0315 (-0.0984, 0.0356, p=3.57E-01) |
| **cg03078551** | -0.0458 (-0.1129, 0.0217, p=1.84E-01) | -0.0278 (-0.0926, 0.0372, p=4.01E-01) | -0.0266 (-0.0916, 0.0385, p=4.23E-01) | -0.0044 (-0.071, 0.0622, p=8.97E-01) | -0.0004 (-0.0674, 0.0666, p=9.91E-01) |
| **cg04927537** | -0.0285 (-0.0958, 0.0391, p=4.09E-01) | -0.0336 (-0.0984, 0.0314, p=3.11E-01) | 0.0031 (-0.062, 0.0682, p=9.25E-01) | 0.0571 (-0.0096, 0.1232, p=9.31E-02) | -0.0037 (-0.0708, 0.0633, p=9.13E-01) |
| **cg07012687** | -0.023 (-0.0903, 0.0446, p=5.05E-01) | -0.0184 (-0.0833, 0.0466, p=5.79E-01) | -0.0082 (-0.0732, 0.057, p=8.06E-01) | 0.0001 (-0.0665, 0.0666, p=9.99E-01) | 0.0295 (-0.0376, 0.0964, p=3.89E-01) |
| **cg08857797** | -0.0174 (-0.0848, 0.0502, p=6.14E-01) | 0.0306 (-0.0344, 0.0954, p=3.56E-01) | 0.0136 (-0.0515, 0.0786, p=6.82E-01) | 0.0154 (-0.0512, 0.0819, p=6.50E-01) | 0.0379 (-0.0292, 0.1047, p=2.68E-01) |
| **cg09664445** | -0.0222 (-0.0895, 0.0454, p=5.20E-01) | -0.0657 (-0.1301, -0.0008, p=4.74E-02) | -0.0574 (-0.122, 0.0078, p=8.42E-02) | 0.0196 (-0.047, 0.0861, p=5.64E-01) | 0.0524 (-0.0147, 0.1191, p=1.25E-01) |
| **cg10508317** | 0.0172 (-0.0503, 0.0846, p=6.18E-01) | 0.0214 (-0.0436, 0.0863, p=5.18E-01) | -0.0505 (-0.1152, 0.0146, p=1.28E-01) | 0.0206 (-0.046, 0.0871, p=5.44E-01) | 0.0359 (-0.0312, 0.1027, p=2.95E-01) |
| **cg11024682** | -0.0804 (-0.1471, -0.013, p=1.95E-02) | -0.0499 (-0.1145, 0.0151, p=1.32E-01) | -0.0061 (-0.0711, 0.059, p=8.55E-01) | -0.0204 (-0.0869, 0.0462, p=5.49E-01) | 0.0219 (-0.0452, 0.0889, p=5.22E-01) |
| **cg11202345** | -0.0162 (-0.0836, 0.0513, p=6.37E-01) | -0.045 (-0.1096, 0.02, p=1.75E-01) | -0.0793 (-0.1436, -0.0142, p=1.70E-02) | 0.0521 (-0.0145, 0.1183, p=1.25E-01) | 0.0136 (-0.0534, 0.0806, p=6.90E-01) |
| **cg13274938** | 0.0113 (-0.0562, 0.0787, p=7.43E-01) | -0.0123 (-0.0772, 0.0527, p=7.11E-01) | 0.003 (-0.0621, 0.068, p=9.29E-01) | 0.0446 (-0.0221, 0.1108, p=1.90E-01) | -0.0571 (-0.1237, 0.01, p=9.52E-02) |
| **cg14509967** | -0.0167 (-0.0841, 0.0508, p=6.28E-01) | -0.0144 (-0.0792, 0.0506, p=6.65E-01) | 0.0316 (-0.0336, 0.0965, p=3.42E-01) | 0.0394 (-0.0273, 0.1057, p=2.47E-01) | 0.0004 (-0.0666, 0.0675, p=9.90E-01) |
| **cg14870271** | -0.0239 (-0.0913, 0.0436, p=4.87E-01) | -0.0508 (-0.1153, 0.0142, p=1.26E-01) | -0.061 (-0.1256, 0.0041, p=6.64E-02) | 0.0197 (-0.047, 0.0862, p=5.63E-01) | -0.0187 (-0.0857, 0.0484, p=5.84E-01) |
| **cg16611584** | -0.0282 (-0.0955, 0.0394, p=4.13E-01) | -0.0314 (-0.0962, 0.0336, p=3.44E-01) | -0.0332 (-0.0981, 0.032, p=3.18E-01) | -0.0113 (-0.0779, 0.0553, p=7.39E-01) | 0.0204 (-0.0468, 0.0873, p=5.52E-01) |
| **cg17836612** | 0.0018 (-0.0657, 0.0693, p=9.58E-01) | -0.0051 (-0.0701, 0.0598, p=8.77E-01) | 0.0279 (-0.0372, 0.0929, p=4.01E-01) | 0 (-0.0666, 0.0666, p=9.99E-01) | -0.0402 (-0.107, 0.0269, p=2.40E-01) |
| **cg18091083** | 0.065 (-0.0025, 0.1319, p=5.92E-02) | -0.0178 (-0.0826, 0.0473, p=5.92E-01) | 0.0174 (-0.0478, 0.0824, p=6.02E-01) | 0.0523 (-0.0144, 0.1184, p=1.24E-01) | 0.0405 (-0.0266, 0.1073, p=2.37E-01) |
| **cg18181703** | 0.0485 (-0.0191, 0.1156, p=1.59E-01) | -0.0334 (-0.0981, 0.0317, p=3.14E-01) | -0.0252 (-0.0901, 0.04, p=4.49E-01) | 0.0107 (-0.056, 0.0772, p=7.54E-01) | 0.0044 (-0.0627, 0.0714, p=8.98E-01) |
| **cg18772573** | -0.0309 (-0.0982, 0.0367, p=3.70E-01) | 0.0664 (0.0014, 0.1308, p=4.52E-02) | 0.0547 (-0.0104, 0.1194, p=9.96E-02) | 0.0113 (-0.0553, 0.0779, p=7.39E-01) | 0.0159 (-0.0512, 0.0829, p=6.43E-01) |
| **cg22713958** | 0.0372 (-0.0304, 0.1044, p=2.80E-01) | -0.0266 (-0.0914, 0.0384, p=4.23E-01) | 0.0401 (-0.0251, 0.1049, p=2.28E-01) | 0.0333 (-0.0333, 0.0997, p=3.27E-01) | -0.021 (-0.088, 0.0461, p=5.39E-01) |
| **cg25178683** | -0.0012 (-0.0686, 0.0663, p=9.73E-01) | -0.0258 (-0.0906, 0.0392, p=4.36E-01) | 0.0121 (-0.0531, 0.0771, p=7.16E-01) | 0.0308 (-0.0359, 0.0972, p=3.65E-01) | -0.0163 (-0.0832, 0.0508, p=6.34E-01) |
| **cg25649826** | 0.0115 (-0.0561, 0.0789, p=7.39E-01) | -0.0158 (-0.0807, 0.0492, p=6.34E-01) | -0.0022 (-0.0673, 0.0629, p=9.47E-01) | 0.0323 (-0.0343, 0.0987, p=3.42E-01) | -0.0189 (-0.0859, 0.0482, p=5.80E-01) |
| **cg26651978** | -0.0116 (-0.0791, 0.0559, p=7.35E-01) | -0.0235 (-0.0883, 0.0415, p=4.78E-01) | 0.0033 (-0.0618, 0.0684, p=9.20E-01) | -0.0469 (-0.1132, 0.0197, p=1.67E-01) | 0.0159 (-0.0512, 0.0829, p=6.42E-01) |
| **cg27470213** | 0.0164 (-0.0511, 0.0838, p=6.33E-01) | 0.0161 (-0.0489, 0.081, p=6.27E-01) | -0.0199 (-0.0849, 0.0453, p=5.50E-01) | -0.0313 (-0.0977, 0.0353, p=3.57E-01) | 0.0166 (-0.0505, 0.0835, p=6.28E-01) |
| **cg27637521** | 0.0225 (-0.0451, 0.0898, p=5.14E-01) | -0.0073 (-0.0722, 0.0577, p=8.25E-01) | -0.0468 (-0.1116, 0.0183, p=1.59E-01) | 0.0048 (-0.0618, 0.0714, p=8.87E-01) | 0.0342 (-0.0329, 0.101, p=3.18E-01) |
| **cg01751802** | -0.0091 (-0.0765, 0.0584, p=7.92E-01) | -0.0364 (-0.1011, 0.0286, p=2.72E-01) | -0.0143 (-0.0793, 0.0509, p=6.68E-01) | -0.0453 (-0.1116, 0.0213, p=1.82E-01) | -0.0294 (-0.0962, 0.0377, p=3.91E-01) |
| **cg02711608** | -0.0245 (-0.0919, 0.043, p=4.77E-01) | -0.0141 (-0.079, 0.0509, p=6.71E-01) | -0.0124 (-0.0775, 0.0527, p=7.09E-01) | 0.0202 (-0.0464, 0.0867, p=5.52E-01) | 0.0007 (-0.0664, 0.0677, p=9.84E-01) |
| **cg04557677** | 0.0156 (-0.0519, 0.083, p=6.51E-01) | -0.0349 (-0.0996, 0.0301, p=2.93E-01) | -0.0058 (-0.0709, 0.0593, p=8.61E-01) | 0.0037 (-0.0629, 0.0703, p=9.13E-01) | 0.0408 (-0.0263, 0.1076, p=2.33E-01) |
| **cg07573872** | 0.0002 (-0.0673, 0.0677, p=9.96E-01) | 0.0129 (-0.0521, 0.0778, p=6.98E-01) | -0.0187 (-0.0837, 0.0464, p=5.73E-01) | -0.0173 (-0.0838, 0.0493, p=6.10E-01) | -0.001 (-0.0681, 0.066, p=9.76E-01) |
| **cg21766592** | 0.0097 (-0.0578, 0.0771, p=7.78E-01) | 0.0144 (-0.0506, 0.0792, p=6.65E-01) | -0.0709 (-0.1354, -0.0059, p=3.27E-02) | -0.0319 (-0.0983, 0.0348, p=3.48E-01) | -0.0143 (-0.0813, 0.0528, p=6.76E-01) |
| **cg22950899** | -0.0293 (-0.0965, 0.0383, p=3.96E-01) | 0.0264 (-0.0386, 0.0912, p=4.25E-01) | 0.0017 (-0.0634, 0.0668, p=9.59E-01) | 0.0286 (-0.0381, 0.095, p=4.00E-01) | -0.0167 (-0.0837, 0.0504, p=6.25E-01) |
| **cg26470501** | 0.0037 (-0.0638, 0.0711, p=9.15E-01) | -0.0554 (-0.1199, 0.0096, p=9.50E-02) | -0.0196 (-0.0846, 0.0455, p=5.55E-01) | -0.035 (-0.1013, 0.0317, p=3.04E-01) | 0.0038 (-0.0633, 0.0708, p=9.13E-01) |
| **cg26950531** | 0.0615 (-0.006, 0.1284, p=7.43E-02) | 0.0291 (-0.0359, 0.0939, p=3.80E-01) | 0.0439 (-0.0213, 0.1087, p=1.87E-01) | -0.0154 (-0.0819, 0.0512, p=6.51E-01) | 0.0247 (-0.0424, 0.0916, p=4.70E-01) |
| **cg03218374** | -0.0145 (-0.0819, 0.053, p=6.74E-01) | -0.0025 (-0.0675, 0.0624, p=9.39E-01) | 0.0329 (-0.0322, 0.0978, p=3.22E-01) | -0.0085 (-0.075, 0.0581, p=8.03E-01) | -0.001 (-0.068, 0.0661, p=9.77E-01) |
| **cg00222799** | 0.01 (-0.0576, 0.0774, p=7.72E-01) | -0.0527 (-0.1172, 0.0123, p=1.12E-01) | -0.0218 (-0.0867, 0.0434, p=5.13E-01) | -0.0428 (-0.1091, 0.0239, p=2.08E-01) | -0.0087 (-0.0757, 0.0584, p=7.99E-01) |
| **cg01881899** | 0.0061 (-0.0614, 0.0736, p=8.59E-01) | -0.0292 (-0.094, 0.0358, p=3.79E-01) | 0.0021 (-0.0631, 0.0671, p=9.51E-01) | 0.0193 (-0.0474, 0.0858, p=5.71E-01) | 0.0591 (-0.0079, 0.1257, p=8.40E-02) |
| **cg06500161** | 0.1083 (0.0411, 0.1745, p=1.63E-03) | -0.066 (-0.1304, -0.0011, p=4.64E-02) | -0.0204 (-0.0854, 0.0448, p=5.40E-01) | 0.0314 (-0.0352, 0.0978, p=3.55E-01) | -0.0427 (-0.1095, 0.0244, p=2.12E-01) |
| **cg10192877** | 0.0115 (-0.056, 0.0789, p=7.38E-01) | 0.0014 (-0.0636, 0.0663, p=9.66E-01) | 0.0509 (-0.0142, 0.1156, p=1.26E-01) | -0.0241 (-0.0905, 0.0426, p=4.79E-01) | 0.004 (-0.0631, 0.071, p=9.08E-01) |
| **cg27243685** | 0.0699 (0.0025, 0.1368, p=4.22E-02) | 0.0208 (-0.0442, 0.0856, p=5.31E-01) | -0.0111 (-0.0762, 0.054, p=7.37E-01) | 0.0087 (-0.0579, 0.0752, p=7.98E-01) | -0.0386 (-0.1054, 0.0285, p=2.60E-01) |
| **cg03682690** | 0.0184 (-0.0491, 0.0858, p=5.93E-01) | -0.0293 (-0.0941, 0.0357, p=3.77E-01) | 0.0067 (-0.0584, 0.0718, p=8.40E-01) | -0.0159 (-0.0824, 0.0508, p=6.41E-01) | -0.0031 (-0.0701, 0.064, p=9.28E-01) |
| **cg06397161** | 0.0241 (-0.0434, 0.0915, p=4.84E-01) | 0.0036 (-0.0613, 0.0686, p=9.13E-01) | -0.0257 (-0.0906, 0.0395, p=4.40E-01) | -0.003 (-0.0696, 0.0636, p=9.29E-01) | 0.0316 (-0.0355, 0.0984, p=3.56E-01) |
| **cg08548559** | -0.0236 (-0.091, 0.0439, p=4.93E-01) | -0.048 (-0.1126, 0.017, p=1.48E-01) | -0.0386 (-0.1034, 0.0266, p=2.46E-01) | -0.0135 (-0.08, 0.0532, p=6.92E-01) | -0.0042 (-0.0712, 0.0629, p=9.03E-01) |
| **cg09182678** | 0.0179 (-0.0497, 0.0852, p=6.04E-01) | -0.0024 (-0.0673, 0.0626, p=9.43E-01) | -0.0102 (-0.0753, 0.0549, p=7.59E-01) | -0.0317 (-0.098, 0.035, p=3.52E-01) | 0.0378 (-0.0293, 0.1046, p=2.70E-01) |
| **cg09349128** | -0.0279 (-0.0952, 0.0396, p=4.17E-01) | -0.048 (-0.1126, 0.017, p=1.47E-01) | -0.0279 (-0.0928, 0.0373, p=4.02E-01) | -0.0357 (-0.1021, 0.0309, p=2.93E-01) | 0.034 (-0.0331, 0.1009, p=3.20E-01) |
| **cg14780837** | 0.0128 (-0.0548, 0.0802, p=7.11E-01) | 0.0359 (-0.0291, 0.1006, p=2.79E-01) | 0.0209 (-0.0442, 0.0859, p=5.29E-01) | 0.007 (-0.0596, 0.0735, p=8.38E-01) | -0.0143 (-0.0813, 0.0528, p=6.77E-01) |
| **cg17194270** | 0.0215 (-0.046, 0.0889, p=5.32E-01) | -0.039 (-0.1037, 0.026, p=2.39E-01) | 0.007 (-0.0581, 0.0721, p=8.32E-01) | -0.0004 (-0.067, 0.0662, p=9.91E-01) | 0.0605 (-0.0065, 0.1271, p=7.69E-02) |
| **cg20496314** | -0.0011 (-0.0686, 0.0664, p=9.75E-01) | -0.0187 (-0.0835, 0.0463, p=5.74E-01) | -0.0402 (-0.105, 0.025, p=2.27E-01) | -0.0145 (-0.081, 0.0521, p=6.70E-01) | 0.0233 (-0.0438, 0.0902, p=4.96E-01) |
| **cg22650271** | -0.007 (-0.0744, 0.0605, p=8.39E-01) | 0.0259 (-0.0391, 0.0907, p=4.35E-01) | 0.0196 (-0.0455, 0.0846, p=5.55E-01) | -0.0632 (-0.1292, 0.0034, p=6.29E-02) | 0.01 (-0.0571, 0.077, p=7.70E-01) |
| **cg27115863** | 0.0126 (-0.0549, 0.08, p=7.15E-01) | -0.0106 (-0.0755, 0.0544, p=7.49E-01) | -0.0482 (-0.113, 0.0169, p=1.47E-01) | -0.0158 (-0.0822, 0.0509, p=6.43E-01) | 0.0167 (-0.0504, 0.0837, p=6.25E-01) |

**Supplementary table 3.** Table of Mendelian Randomisation type analyses, using a mixed model combining mother and child data, to further assess whether there is a causal association from BMI to each of the 135 CpG sites used to construct the methylation score.

| **CpG** | **Estimate** | **P-value** | **Lower CI** | **Upper CI** |
| --- | --- | --- | --- | --- |
| **cg01455178** | -0.00390 | 6.76E-01 | -0.02220 | 0.01439 |
| **cg03725309** | 0.00061 | 9.27E-01 | -0.01241 | 0.01364 |
| **cg08639339** | -0.00533 | 3.85E-01 | -0.01733 | 0.00667 |
| **cg10092518** | -0.00616 | 3.25E-01 | -0.01839 | 0.00607 |
| **cg10717869** | 0.00017 | 9.78E-01 | -0.01198 | 0.01234 |
| **cg11673687** | -0.00130 | 8.27E-01 | -0.01287 | 0.01028 |
| **cg12458003** | 0.00297 | 7.00E-01 | -0.01211 | 0.01806 |
| **cg12484113** | 0.00275 | 6.44E-01 | -0.00888 | 0.01437 |
| **cg12593793** | -0.00698 | 2.28E-01 | -0.01828 | 0.00433 |
| **cg14476101** | -0.00375 | 7.43E-01 | -0.02608 | 0.01861 |
| **cg17901584** | -0.00624 | 5.71E-01 | -0.02777 | 0.01528 |
| **cg23172671** | 0.01335 | 1.71E-01 | -0.00571 | 0.03242 |
| **cg23998749** | 0.00667 | 3.59E-01 | -0.00753 | 0.02086 |
| **cg24145109** | 0.00223 | 8.46E-01 | -0.02016 | 0.02461 |
| **cg24678869** | -0.00002 | 9.98E-01 | -0.01023 | 0.01020 |
| **cg25217710** | -0.00156 | 8.33E-01 | -0.01608 | 0.01296 |
| **cg04011474** | -0.01127 | 1.22E-01 | -0.02551 | 0.00297 |
| **cg04286697** | 0.00016 | 9.82E-01 | -0.01439 | 0.01473 |
| **cg06876354** | 0.00603 | 2.63E-01 | -0.00451 | 0.01657 |
| **cg12001357** | -0.00716 | 4.14E-01 | -0.02431 | 0.00999 |
| **cg13139542** | 0.00012 | 9.83E-01 | -0.01097 | 0.01123 |
| **cg14017402** | 0.00358 | 7.40E-01 | -0.01755 | 0.02469 |
| **cg19017142** | -0.01445 | 9.99E-02 | -0.03161 | 0.00271 |
| **cg26191447** | 0.01445 | 1.58E-01 | -0.00558 | 0.03448 |
| **cg00108715** | 0.00087 | 8.98E-01 | -0.01237 | 0.01410 |
| **cg01368219** | 0.01293 | 1.04E-01 | -0.00263 | 0.02848 |
| **cg01526748** | 0.02242 | 5.38E-02 | -0.00033 | 0.04520 |
| **cg01671681** | -0.01540 | 3.96E-02 | -0.03002 | -0.00077 |
| **cg07730360** | 0.01204 | 1.75E-01 | -0.00536 | 0.02946 |
| **cg17641710** | -0.00162 | 8.04E-01 | -0.01443 | 0.01118 |
| **cg18098839** | 0.00026 | 9.61E-01 | -0.00992 | 0.01042 |
| **cg22012981** | 0.00514 | 4.10E-01 | -0.00708 | 0.01738 |
| **cg05119988** | -0.00336 | 7.70E-01 | -0.02579 | 0.01907 |
| **cg06690548** | 0.00389 | 5.46E-01 | -0.00870 | 0.01649 |
| **cg07094298** | -0.00339 | 7.76E-01 | -0.02672 | 0.01993 |
| **cg02286155** | 0.00715 | 2.15E-01 | -0.00413 | 0.01844 |
| **cg04483863** | 0.00656 | 3.57E-01 | -0.00738 | 0.02052 |
| **cg10179300** | 0.00804 | 1.38E-01 | -0.00256 | 0.01863 |
| **cg13276570** | 0.00408 | 4.03E-01 | -0.00546 | 0.01361 |
| **cg13305415** | 0.00343 | 7.55E-01 | -0.01808 | 0.02493 |
| **cg26403843** | -0.01883 | 8.34E-02 | -0.04009 | 0.00243 |
| **cg13123009** | -0.00268 | 6.44E-01 | -0.01400 | 0.00865 |
| **cg14352682** | -0.00029 | 9.59E-01 | -0.01147 | 0.01089 |
| **cg17501210** | -0.01144 | 1.51E-01 | -0.02703 | 0.00414 |
| **cg17738521** | -0.00007 | 9.89E-01 | -0.01030 | 0.01016 |
| **cg22875823** | 0.01135 | 2.49E-01 | -0.00792 | 0.03060 |
| **cg04816311** | -0.00161 | 9.23E-01 | -0.03406 | 0.03084 |
| **cg13134297** | -0.00790 | 4.46E-01 | -0.02816 | 0.01237 |
| **cg21429551** | -0.00789 | 5.77E-01 | -0.03564 | 0.01981 |
| **cg02571142** | -0.00900 | 3.95E-01 | -0.02974 | 0.01171 |
| **cg07960624** | -0.01633 | 1.86E-01 | -0.04048 | 0.00780 |
| **cg24531955** | -0.00610 | 4.94E-01 | -0.02354 | 0.01133 |
| **cg25392060** | -0.00596 | 4.15E-01 | -0.02026 | 0.00833 |
| **cg26361535** | 0.01033 | 2.43E-01 | -0.00696 | 0.02764 |
| **cg02716826** | -0.00073 | 9.09E-01 | -0.01329 | 0.01182 |
| **cg07504977** | -0.01164 | 2.40E-01 | -0.03099 | 0.00772 |
| **cg09491962** | -0.00391 | 5.34E-01 | -0.01620 | 0.00837 |
| **cg14333542** | -0.00662 | 2.77E-01 | -0.01859 | 0.00535 |
| **cg15880704** | -0.00229 | 6.64E-01 | -0.01261 | 0.00803 |
| **cg17782974** | -0.00650 | 5.04E-01 | -0.02550 | 0.01250 |
| **cg26955383** | 0.00456 | 4.93E-01 | -0.00844 | 0.01756 |
| **cg00574958** | 0.00133 | 7.38E-01 | -0.00649 | 0.00915 |
| **cg11152384** | -0.00720 | 1.79E-01 | -0.01767 | 0.00327 |
| **cg11261850** | -0.00531 | 6.53E-01 | -0.02843 | 0.01784 |
| **cg13028635** | 0.00929 | 1.59E-01 | -0.00361 | 0.02220 |
| **cg17058475** | 0.00025 | 9.58E-01 | -0.00904 | 0.00953 |
| **cg26800893** | 0.00047 | 9.00E-01 | -0.00690 | 0.00784 |
| **cg26894079** | 0.00475 | 6.73E-01 | -0.01727 | 0.02677 |
| **cg13708645** | -0.00091 | 9.14E-01 | -0.01737 | 0.01558 |
| **cg10474597** | 0.00356 | 6.88E-01 | -0.01373 | 0.02084 |
| **cg19750657** | 0.00288 | 7.06E-01 | -0.01206 | 0.01783 |
| **cg10919522** | -0.00512 | 5.88E-01 | -0.02360 | 0.01335 |
| **cg27394566** | 0.00404 | 5.30E-01 | -0.00859 | 0.01669 |
| **cg06192883** | 0.00436 | 4.50E-01 | -0.00694 | 0.01567 |
| **cg07037944** | -0.00284 | 6.25E-01 | -0.01420 | 0.00852 |
| **cg07814318** | 0.00128 | 9.14E-01 | -0.02204 | 0.02461 |
| **cg20507228** | -0.00783 | 5.88E-01 | -0.03606 | 0.02040 |
| **cg21670987** | -0.00529 | 6.07E-01 | -0.02544 | 0.01485 |
| **cg01243823** | -0.02088 | 2.13E-02 | -0.03859 | -0.00317 |
| **cg03500056** | 0.00236 | 7.25E-01 | -0.01072 | 0.01544 |
| **cg06946797** | -0.00954 | 2.32E-01 | -0.02514 | 0.00606 |
| **cg07021906** | 0.01509 | 1.19E-01 | -0.00382 | 0.03399 |
| **cg07955474** | -0.00708 | 3.92E-01 | -0.02328 | 0.00911 |
| **cg09607047** | -0.00176 | 8.40E-01 | -0.01884 | 0.01532 |
| **cg16739178** | 0.00182 | 8.34E-01 | -0.01517 | 0.01881 |
| **cg23813257** | 0.00030 | 9.59E-01 | -0.01092 | 0.01150 |
| **cg01130991** | 0.00100 | 9.31E-01 | -0.02138 | 0.02337 |
| **cg01597398** | -0.00783 | 2.26E-01 | -0.02047 | 0.00481 |
| **cg01798813** | 0.00017 | 9.72E-01 | -0.00929 | 0.00963 |
| **cg03078551** | -0.00953 | 1.56E-01 | -0.02268 | 0.00361 |
| **cg04927537** | -0.00049 | 9.59E-01 | -0.01919 | 0.01819 |
| **cg07012687** | -0.00002 | 9.98E-01 | -0.01667 | 0.01664 |
| **cg08857797** | 0.01159 | 2.33E-01 | -0.00739 | 0.03057 |
| **cg09664445** | 0.00137 | 8.55E-01 | -0.01330 | 0.01605 |
| **cg10508317** | -0.00676 | 1.32E-01 | -0.01554 | 0.00202 |
| **cg11024682** | -0.00405 | 3.87E-01 | -0.01320 | 0.00510 |
| **cg11202345** | -0.00830 | 2.76E-01 | -0.02322 | 0.00660 |
| **cg13274938** | 0.00989 | 2.33E-01 | -0.00630 | 0.02608 |
| **cg14509967** | 0.00885 | 2.38E-01 | -0.00584 | 0.02353 |
| **cg14870271** | -0.01487 | 6.21E-02 | -0.03046 | 0.00071 |
| **cg16611584** | -0.00972 | 5.32E-01 | -0.04017 | 0.02076 |
| **cg17836612** | -0.00101 | 8.99E-01 | -0.01656 | 0.01454 |
| **cg18091083** | 0.01943 | 1.98E-01 | -0.01006 | 0.04891 |
| **cg18181703** | -0.00783 | 3.68E-01 | -0.02484 | 0.00919 |
| **cg18772573** | 0.01229 | 1.94E-01 | -0.00620 | 0.03077 |
| **cg22713958** | 0.00853 | 2.40E-01 | -0.00568 | 0.02275 |
| **cg25178683** | -0.00317 | 7.29E-01 | -0.02111 | 0.01476 |
| **cg25649826** | 0.00106 | 8.72E-01 | -0.01175 | 0.01385 |
| **cg26651978** | -0.00522 | 5.27E-01 | -0.02139 | 0.01096 |
| **cg27470213** | -0.01110 | 1.91E-01 | -0.02771 | 0.00553 |
| **cg27637521** | -0.00501 | 1.73E-01 | -0.01216 | 0.00216 |
| **cg01751802** | -0.01155 | 1.83E-01 | -0.02851 | 0.00541 |
| **cg02711608** | -0.00006 | 9.91E-01 | -0.01039 | 0.01026 |
| **cg04557677** | -0.00102 | 7.38E-01 | -0.00700 | 0.00495 |
| **cg07573872** | -0.01022 | 2.89E-01 | -0.02906 | 0.00861 |
| **cg21766592** | -0.01013 | 6.59E-02 | -0.02088 | 0.00064 |
| **cg22950899** | 0.00489 | 5.74E-01 | -0.01212 | 0.02190 |
| **cg26470501** | -0.00822 | 1.61E-01 | -0.01970 | 0.00326 |
| **cg26950531** | 0.00476 | 7.27E-01 | -0.02196 | 0.03149 |
| **cg03218374** | 0.00465 | 5.13E-01 | -0.00925 | 0.01855 |
| **cg00222799** | -0.01069 | 1.99E-01 | -0.02695 | 0.00558 |
| **cg01881899** | -0.00413 | 3.42E-01 | -0.01264 | 0.00437 |
| **cg06500161** | 0.00238 | 7.47E-01 | -0.01204 | 0.01680 |
| **cg10192877** | 0.00388 | 5.12E-01 | -0.00770 | 0.01548 |
| **cg27243685** | -0.00104 | 8.95E-01 | -0.01644 | 0.01437 |
| **cg03682690** | 0.00205 | 7.52E-01 | -0.01059 | 0.01469 |
| **cg06397161** | 0.00022 | 9.73E-01 | -0.01286 | 0.01332 |
| **cg08548559** | -0.01729 | 4.72E-02 | -0.03432 | -0.00026 |
| **cg09182678** | -0.00587 | 2.54E-01 | -0.01594 | 0.00419 |
| **cg09349128** | -0.00449 | 4.83E-01 | -0.01705 | 0.00805 |
| **cg14780837** | 0.00318 | 6.75E-01 | -0.01167 | 0.01803 |
| **cg17194270** | 0.00413 | 7.08E-01 | -0.01742 | 0.02568 |
| **cg20496314** | -0.01103 | 1.84E-01 | -0.02726 | 0.00520 |
| **cg22650271** | -0.00388 | 5.45E-01 | -0.01645 | 0.00871 |
| **cg27115863** | -0.01186 | 1.56E-01 | -0.02823 | 0.00450 |

**Supplementary table 4.** Results from the Mendelian Randomisation type analyses for the direction of methylation variation causing BMI.

| **CpG** | **SNP** | **Proxy SNP** | **Beta** | **SE** | **N** | **p-value** |
| --- | --- | --- | --- | --- | --- | --- |
| **cg25392060** | rs12677618 | NA | -7.00E-04 | 0.0041 | 235522 | 0.86 |
| **cg01455178** | rs6673687 | NA | 0.0103 | 0.0037 | 235727 | 0.005 |
| **cg21670987** | rs7172615 | NA | 0.0083 | 0.0043 | 236092 | 0.05 |
| **cg25649826** | rs7215369 | NA | -0.001 | 0.0045 | 230313 | 0.82 |
| **cg21670987** | rs12903325 | NA | 0.0075 | 0.0042 | 235743 | 0.07 |
| **cg13708645** | rs1549349 | NA | -0.0045 | 0.0037 | 335771 | 0.22 |
| **cg01751802** | rs11878417 | NA | -8.00E-04 | 0.0037 | 235862 | 0.83 |
| **cg07955474** | rs16882 | NA | -9.00E-04 | 0.0047 | 234763 | 0.85 |
| **cg23813257** | rs1064948 | NA | -0.0054 | 0.0041 | 228964 | 0.19 |
| **cg01751802** | rs6511727 | NA | -0.0011 | 0.003 | 338998 | 0.71 |
| **cg04286697** | rs2595507 | NA | -0.0084 | 0.0045 | 235685 | 0.06 |
| **cg22950899** | rs2058110 | NA | -0.0021 | 0.0048 | 188352 | 0.66 |
| **cg11152384** | rs230539 | NA | -0.0034 | 0.0039 | 235792 | 0.38 |
| **cg12001357** | rs2099489 | NA | 4.00E-04 | 0.0061 | 179315 | 0.95 |
| **cg14017402** | rs2278536 | NA | -0.0061 | 0.0042 | 181293 | 0.15 |
| **cg23813257** | rs13335800 | NA | -0.0054 | 0.004 | 224535 | 0.18 |
| **cg07955474** | rs11646550 | NA | -0.0019 | 0.0053 | 228893 | 0.72 |
| **cg03500056** | rs1592457 | NA | 0.0031 | 0.0041 | 224078 | 0.45 |
| **cg08548559** | rs9621221 | NA | -0.0076 | 0.0036 | 236207 | 0.03 |
| **cg01751802** | rs7252965 | rs3745682 | 8.00E-04 | 0.0037 | 236028 | 0.83 |
| **cg01751802** | rs10421221 | rs4804574 | -8.00E-04 | 0.0037 | 236081 | 0.83 |
| **cg04286697** | rs2595503 | rs2290130 | -0.0087 | 0.0042 | 236094 | 0.04 |
| **cg07955474** | rs1519978 | rs16882 | -9.00E-04 | 0.0047 | 234763 | 0.85 |
| **cg08548559** | rs5997938 | rs4820961 | -0.0076 | 0.0036 | 236196 | 0.03 |
| **cg08548559** | rs230497 | rs230498 | -0.0053 | 0.0039 | 235825 | 0.17 |
| **cg08548559** | rs8141987 | rs9621216 | -0.0075 | 0.0036 | 236213 | 0.04 |
| **cg12001357** | rs142273229 | rs2099489 | 4.00E-04 | 0.0061 | 179315 | 0.95 |
| **cg14017402** | rs10178540 | rs2278536 | -0.0061 | 0.0042 | 181293 | 0.15 |
| **cg24531955** | rs6557652 | rs11135720 | 0.0014 | 0.0036 | 235962 | 0.70 |
| **cg25649826** | rs4985973 | rs4985974 | -0.0015 | 0.0045 | 236006 | 0.74 |
| **cg26403843** | rs7732652 | rs1473247 | 0.0017 | 0.0033 | 339062 | 0.61 |
| **cg26403843** | rs10077858 | rs1897565 | -3.00E-04 | 0.0041 | 236045 | 0.94 |
| **cg26470501** | rs56702353 | rs17728272 | 0.0068 | 0.0046 | 222495 | 0.14 |

**Supplementary table 5.** Confounders analysis (mother’s smoking and education level) results from 7 different linear models with methylation score as the dependent variable and different combinations of BMI, mother’s smoking and mother’s education as the independent variables, across the 5 time points.

|  |  | **Beta value from Linear model (CI), p-value** | | | | |
| --- | --- | --- | --- | --- | --- | --- |
| **Linear Model** |  | **Birth** | **Childhood** | **Adolescence** | **Pregnancy** | **Middle-age** |
| **Methylation ~ BMI** | BMI | 0.0003 (0.0002, 0.0005), 9.79E-06^a,b^ | 0.05 (0.01, 0.08), 0.005^a,b^ | 0.05 (0.03, 0.07), 1.05E-06^a,b^ | 0.04 (0.02, 0.06), 2.72E-05^a,b^ | 0.06 (0.05, 0.07), 2.15E-21^a,b^ |
| **Methylation ~ BMI + Mother smokes** | BMI | 0.0003 (0.0001, 0.0004), 0.0001^a,b^ | 0.05 (0.01, 0.08), 0.008^a^ | 0.05 (0.03, 0.07), 3.46E-06^a,b^ | 0.04 (0.02, 0.07), 5.92E-04^a,b^ | 0.06 (0.04, 0.07), 132E-14^a,b^ |
|  | Mother smokes | -0.18 (-0.35, -0.007), 0.04^a^ | -0.001 (-0.17, 0.17), 0.99 | 0.05 (-0.13, 0.24), 0.57 | 0.03 (-0.19, 0.25), 0.79 | 0.11 (-0.10, 0.31), 0.30 |
| **Methylation ~ Mother smokes** | Mother smokes | -0.20 (-0.37, -0.02), 0.03^a^ | 0.02 (-0.15, 0.18), 0.83 | 0.09 (-0.09, 0.27), 0.34 | 0.07 (-0.16, 0.29), 0.55 | 0.15 (-0.06, 0.37), 0.16 |
| **Methylation ~ BMI + Education of mother** | BMI | 0.0003 (0.0002, 0.0005), 2.51E-05^a,b^ | 0.05 (0.01, 0.08), 0.005^a,b^ | 0.05 (0.03, 0.07), 1.70E-06^a,b^ | 0.04 (0.02, 0.06), 4.92E-05^a,b^ | 0.06 (0.05, 0.07), 4.94E-20^a,b^ |
|  | Education (Vocational) | -0.03 (-0.39, 0.33), 0.88 | 0.15 (-0.21, 0.52), 0.42 | 0.15 (-0.24, 0.53), 0.45 | 0.09 (-0.29, 0.47), 0.64 | 0.32 (-06, 0.69), 0.10 |
|  | Education (O-level) | 0.05 (-0.24, 0.34), 0.76 | -0.009 (-0.30, 0.28), 0.95 | 0.11 (-0.20, 0.43), 0.48 | 0.15 (-0.16, 0.46), 0.34 | 0.14 (-0.17, 0.45), 0.37 |
|  | Education (A-level) | 0.12 (-0.18, 0.41), 0.44 | 0.003 (-0.29, 0.30), 0.98 | 0.09 (-0.23, 0.41), 0.58 | 0.04 (-0.27, 0.35), 0.78 | 0.16 (-0.15, 0.47), 0.32 |
|  | Education (Degree) | 0.08 (-0.23, 0.38), 0.62 | 0.12 (-0.19, 0.42), 0.45 | 0.03 (-0.29, 0.36), 0.84 | 0.14 (-0.19, 0.46), 0.41 | 0.18 (-0.14, 0.51), 0.28 |
| **Methylation ~ BMI + Mother smokes + Education of mother** | BMI | 0.0003 (0.0001, 0.0004), 0.0002^a,b^ | 0.05 (0.01, 0.08), 0.009^a^ | 0.05 (0.03, 0.07), 4.15E-06^a,b^ | 0.04 (0.02, 0.07), 5.31E-04^a,b^ | 0.06 (0.04, 0.07), 1.52E-14^a,b^ |
|  | Mother smokes | -0.14 (-0.32, 0.04), 0.12 | 0.03 (0.01, 0.08), 0.73 | 0.07 (-0.12, 0.26), 0.46 | 0.02 (-0.22, 0.25), 0.89 | 0.11 (-0.10, 0.32), 0.29 |
|  | Education (Vocational) | -0.05 (-0.42, 0.33), 0.80 | 0.18 (-0.20, 0.55), 0.36 | 0.18 (-0.22, 0.58), 0.39 | 0.24 (-0.27, 0.75), 0.36 | 0.33 (-0.15, 0.80), 0.18 |
|  | Education (O-level) | -0.006 (-0.31, 0.30), 0.97 | -0.03 (-0.33, 0.28), 0.87 | 0.14 (-0.19, 0.47), 0.40 | 0.21 (-0.20, 0.61), 0.32 | 0.18 (-0.21, 0.56), 0.38 |
|  | Education (A-level) | 0.07 (-0.24, 0.38), 0.66 | 0.02 (-0.28, 0.33), 0.88 | 0.12 (-0.21, 0.46), 0.47 | 0.10 (-0.31, 0.50), 0.64 | 0.27 (-0.12, 0.66), 0.17 |
|  | Education (Degree) | 0.04 (-0.29, 0.36), 0.82 | 0.14 (-0.18, 0.46), 0.39 | 0.06 (-0.28, 0.41), 0.72 | 0.20 (-0.22, 0.61), 0.35 | 0.31 (-0.09, 0.71), 0.14 |
| **Methylation ~ Education of mother** | Education (Vocational) | -0.02 (-0.39, 0.34), 0.91 | 0.16 (-0.20, 0.53), 0.39 | 0.13 (-0.26, 0.52), 0.52 | 0.12 (-0.27, 0.50), 0.55 | 0.28 (-0.12, 0.68), 0.17 |
|  | Education (O-level) | 0.07 (-0.39, 0.34), 0.66 | -0.009 (-0.30, 0.28), 0.95 | 0.12 (-0.20, 0.44), 0.47 | 0.13 (-0.18, 0.44), 0.41 | 0.01 (-0.31, 0.34), 0.93 |
|  | Education (A-level) | 0.13 (-0.17, 0.43), 0.40 | 0.002 (-0.30, 0.28), 0.99 | 0.06 (-0.26, 0.38), 0.71 | 0.01 (-0.30, 0.33), 0.70 | 0.03 (-0.30, 0.36), 0.86 |
|  | Education (Degree) | 0.10 (-0.21, 0.41), 0.51 | 0.10 (-0.20, 0.40), 0.51 | -0.02 (-0.35, 0.31), 0.90 | 0.07 (-0.25, 0.39), 0.67 | -0.05 (-0.39, 0.29), 0.76 |
| **Methylation ~ Education of mother + Mother smokes** | Mother smokes | -0.15 (-0.34, 0.03), 0.09 | 0.05 (-0.12, 0.22), 0.58 | 0.10 (-0.09, 0.29), 0.29 | 0.05 (-0.18, 0.28), 0.68 | 0.14 (-0.08, 0.37), 0.20 |
|  | Education (Vocational) | -0.04 (-0.42, 0.34), 0.83 | 0.19 (-0.19, 0.57), 0.33 | 0.16 (-0.24, 0.57), 0.43 | 0.25 (-0.26, 0.76), 0.34 | 0.31 (-0.20, 0.81), 0.23 |
|  | Education (O-level) | 0.01 (-0.30, 0.32), 0.95 | -0.03 (-0.33, 0.28), 0.87 | 0.15 (-0.18, 0.48), 0.43 | 0.18 (-0.22, 0.59), 0.38 | 0.06 (-0.35, 0.47), 0.77 |
|  | Education (A-level) | 0.08 (-0.23, 0.39), 0.62 | 0.02 (-0.28, 0.33), 0.88 | 0.10 (-0.23, 0.44), 0.55 | 0.08 (-0.33, 0.49), 0.70 | 0.18 (-0.23, 0.59), 0.39 |
|  | Education (Degree) | 0.06 (-0.27, 0.38), 0.72 | 0.13 (-0.19, 0.45), 0.43 | 0.02 (-0.33, 0.37), 0.91 | 0.13 (-0.29, 0.55), 0.53 | 0.11 (-0.31, 0.53), 0.61 |

a= significant at the 0.05 p-value threshold, b= significant at the adjusted p-value threshold of 0.007 (0.05/7, as there were 7 models).

**Supplementary table 6.** Results from ANOVA tests comparing the model adjusting for methylation score and BMI (outcome ~ BMI + methylation score) with the model with just BMI (outcome ~ BMI) in the first columns and then the full model (outcome ~ methylation score + genetic score) is compared with the model with just methylation score (outcome ~ methylation score) in the other columns. These tests were additionally conducted for the models with and without adjustment for BMI, as indicated in the table. The data used was the adolescent data for children and the middle-aged data for mothers.

|  | **Child (adolescence)** | | | | | | **Mother (middle age)** | | | | | |
| --- | --- | --- | --- | --- | --- | --- | --- | --- | --- | --- | --- | --- |
|  | **Comparing:**  **1.** **outcome ~ BMI**  **2. outcome ~ BMI + methylation score** | | **Comparing:**  **1. outcome ~ methylation score**  **2. outcome ~ methylation score + genetic score** | | **Comparing:**  **1. outcome ~ methylation score + BMI**  **2. outcome ~ methylation score + genetic score + BMI** | | **Comparing:**  **1.** **outcome ~ BMI**  **2. outcome ~ BMI + methylation score** | | **Comparing:**  **1. outcome ~ methylation score**  **2. outcome ~ methylation score + genetic score** | | **Comparing:**  **1. outcome ~ methylation score + BMI**  **2. outcome ~ methylation score + genetic score + BMI** | |
|  | **F** | **p-value** | **F** | **p-value** | **F** | **p-value** | **F** | **p-value** | **F** | **p-value** | **F** | **p-value** |
| **log(triglycerides)** | 1.44 | 0.18 | 0.03 | 0.88 | 1.42 | 0.23 | 2.81 | 3.00E-03 | 1.69 | 0.19 | 8.71 | 3.27E-03 |
| **LDL** | 2.50 | 0.02 | 0.48 | 0.49 | 0.19 | 0.67 | 2.36 | 0.01 | 0.09 | 0.76 | 0.50 | 0.48 |
| **log(glucose)** | 2.20 | 0.03 | 0.65 | 0.42 | 0.01 | 0.92 | 1.73 | 0.08 | 5.05 | 0.02 | 1.71 | 0.19 |
| **log(insulin)** | 0.84 | 0.56 | 0.07 | 0.79 | 2.80 | 0.09 | 1.52 | 0.14 | 0.64 | 0.43 | 1.92 | 0.17 |
| **SBP** | 1.85 | 0.08 | 16.11 | 6.60E-05 | 3.78 | 0.05 | 2.11 | 0.03 | 0.04 | 0.83 | 2.36 | 0.13 |
| **DBP** | 1.14 | 0.33 | 4.63 | 0.03 | 0.17 | 0.68 | 1.44 | 0.17 | 1.45 | 0.23 | 0.24 | 0.62 |

**Supplementary table 7.** Results from ANOVA tests comparing the model adjusting for genetic score and BMI (outcome ~ BMI + genetic score) with the model with just BMI (outcome ~ BMI) in the first columns and then the full model (outcome ~ methylation score + genetic score) with the model with just genetic score (outcome ~ genetic score) in the other columns. These tests were additionally conducted for the models with and without adjustment for BMI, as indicated in the table. The data used was the adolescent data for children and the middle-aged data for mothers.

|  | **Child** | | | | | | **Mother** | | | | | |
| --- | --- | --- | --- | --- | --- | --- | --- | --- | --- | --- | --- | --- |
|  | **Comparing:**  **1.** **outcome ~ BMI**  **2. outcome ~ BMI + genetic score** | | **Comparing:**  **1. outcome ~ genetic score**  **2. outcome ~ methylation score + genetic score** | | **Comparing:**  **1. outcome ~ genetic score + BMI**  **2. outcome ~ methylation score + genetic score + BMI** | | **Comparing:**  **1.** **outcome ~ BMI**  **2. outcome ~ BMI + genetic score** | | **Comparing:**  **1. outcome ~ genetic score**  **2. outcome ~ methylation score + genetic score** | | **Comparing:**  **1. outcome ~ genetic score + BMI**  **2. outcome ~ methylation score + genetic score + BMI** | |
|  | **F** | **p-value** | **F** | **p-value** | **F** | **p-value** | **F** | **p-value** | **F** | **p-value** | **F** | **p-value** |
| **log(triglycerides)** | 1.44 | 0.18 | 2.93 | 5.00E-03 | 1.47 | 0.18 | 3.43 | 2.01E-04 | 7.21 | 4.66E-10 | 2.74 | 3.80E-03 |
| **LDL** | 2.21 | 0.03 | 2.80 | 7.00E-03 | 2.51 | 0.02 | 2.17 | 0.02 | 3.41 | 4.10E-04 | 2.34 | 0.01 |
| **log(glucose)** | 1.92 | 0.05 | 1.89 | 0.07 | 2.18 | 0.03 | 1.73 | 0.07 | 2.31 | 0.01 | 1.71 | 0.08 |
| **log(insulin)** | 1.08 | 0.37 | 2.44 | 0.02 | 0.83 | 0.56 | 1.55 | 0.12 | 6.92 | 1.39E-09 | 1.51 | 0.14 |
| **SBP** | 2.10 | 0.03 | 2.90 | 5.38E-03 | 1,71 | 0.10 | 2.14 | 0.02 | 3.45 | 3.51E-04 | 2.17 | 0.02 |
| **DBP** | 1.02 | 0.42 | 2.15 | 0.04 | 1.14 | 0.34 | 1.32 | 0.22 | 2.01 | 0.04 | 1.46 | 0.16 |

**Supplementary figure 1.** Cross-lagged model for sub model with path from childhood DNA methylation to adolescent BMI dropped, in children.

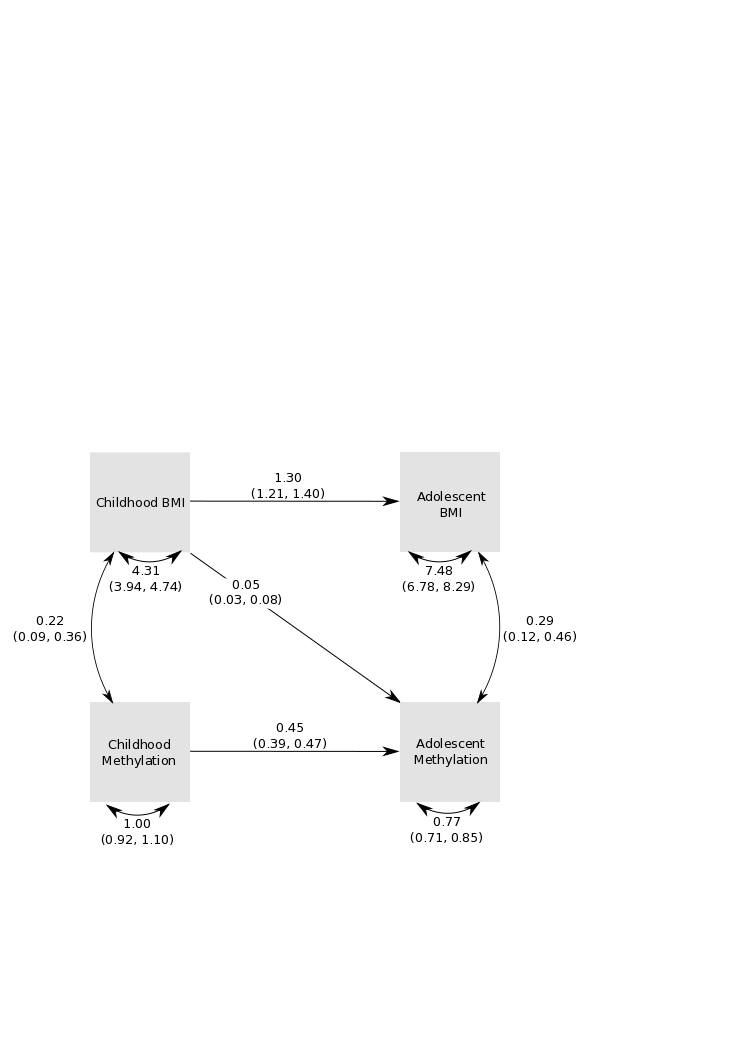

**Supplementary figure 2.** Cross-lagged model for sub model path from DNA methylation score in pregnancy to BMI in middle-age dropped, in mothers.

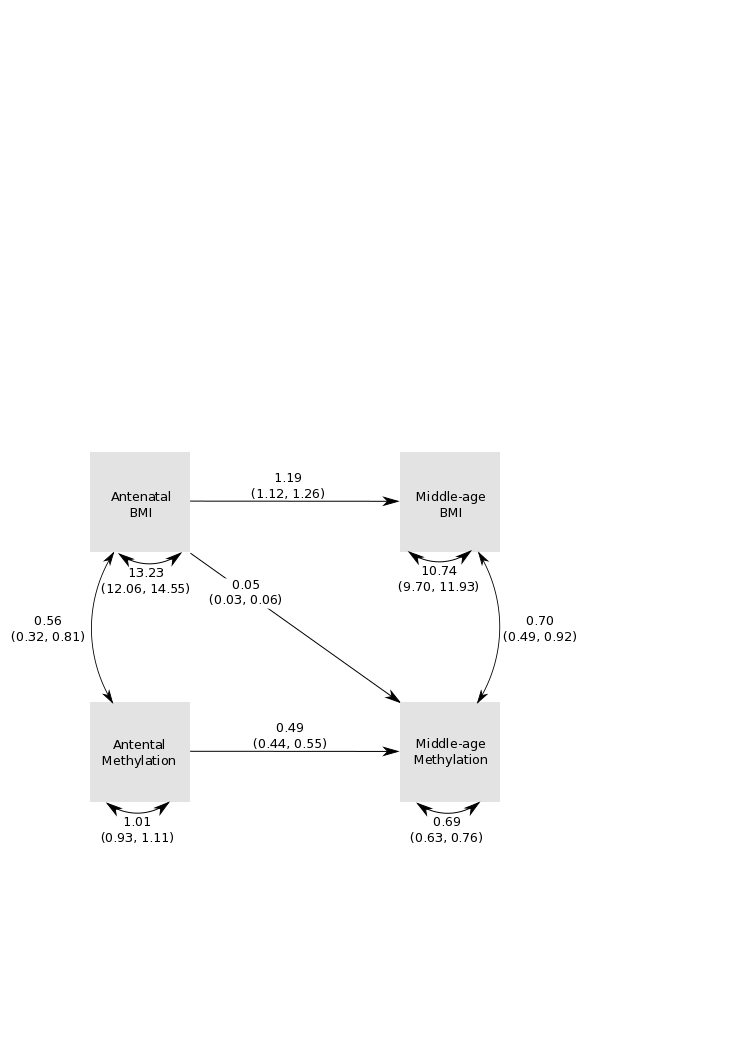
